# Spatial programming of fibroblasts promotes resolution of tissue inflammation through immune cell exclusion

**DOI:** 10.1101/2024.09.20.614064

**Authors:** Patricia Reis Nisa, Christopher Mahony, Annie Hackland, Elizabeth Clay, Paulynn Suyin Chin, Lucy-Jayne Marsh, Chrissy Bolton, Samuel Kemble, Charlotte G. Smith, Georgiana Neag, Ruchir Singh, Jason D. Turner, Andrew Filer, Karim Raza, Jean-Baptiste Richard, Kim S. Midwood, Christopher D. Buckley, Kevin Wei, Helen M. McGettrick, Mark Coles, Adam P. Croft

**Author notes:** These authors contributed equally (Patricia Reis Nisa, Christopher Mahony). Corresponding authors: Professor Adam Paul Croft and Dr Chris Mahony. Rheumatology Research Group, Institute for Inflammation and Ageing, College of Medical and Dental Sciences, University of Birmingham, Queen Elizabeth Hospital, Birmingham, B15 2WD, UK; NIHR Biomedical Research Centre, Institute of Translational Medicine, Birmingham, United Kingdom, B15 2TH.

## Abstract

The role of fibroblasts in determining tissue topography and immune cell organisation within chronically inflamed tissues is poorly understood. Herein, we use multi-omic spatial analysis to define the cellular zonation pattern of the synovium in patients with inflammatory arthritis, identifying discrete tissue niches underpinned by spatially programmed subsets of synovial fibroblasts. We observe that perivascular fibroblasts switch on distinct matrix programs in response to cytokine signalling from neighbouring cells, forming adapted tissue niches that either permit or restrict immune cell trafficking. Specifically, IFN-γ−responsive fibroblasts form a pathogenic lymphocyte-permissive niche that supports the persistence of leukocytes in the tissue, whilst TGF-β−responsive, matrix-synthesising fibroblasts comprise a reparative niche, composed of a collagen-rich barrier around blood vessels that restricts leukocyte migration and promotes resolution of tissue inflammation. Augmentation of such endogenous pathways to promote resolution of inflammation may offer therapeutically tractable approaches for restoration of tissue homeostasis.

GRAPHICAL ABSTRACT

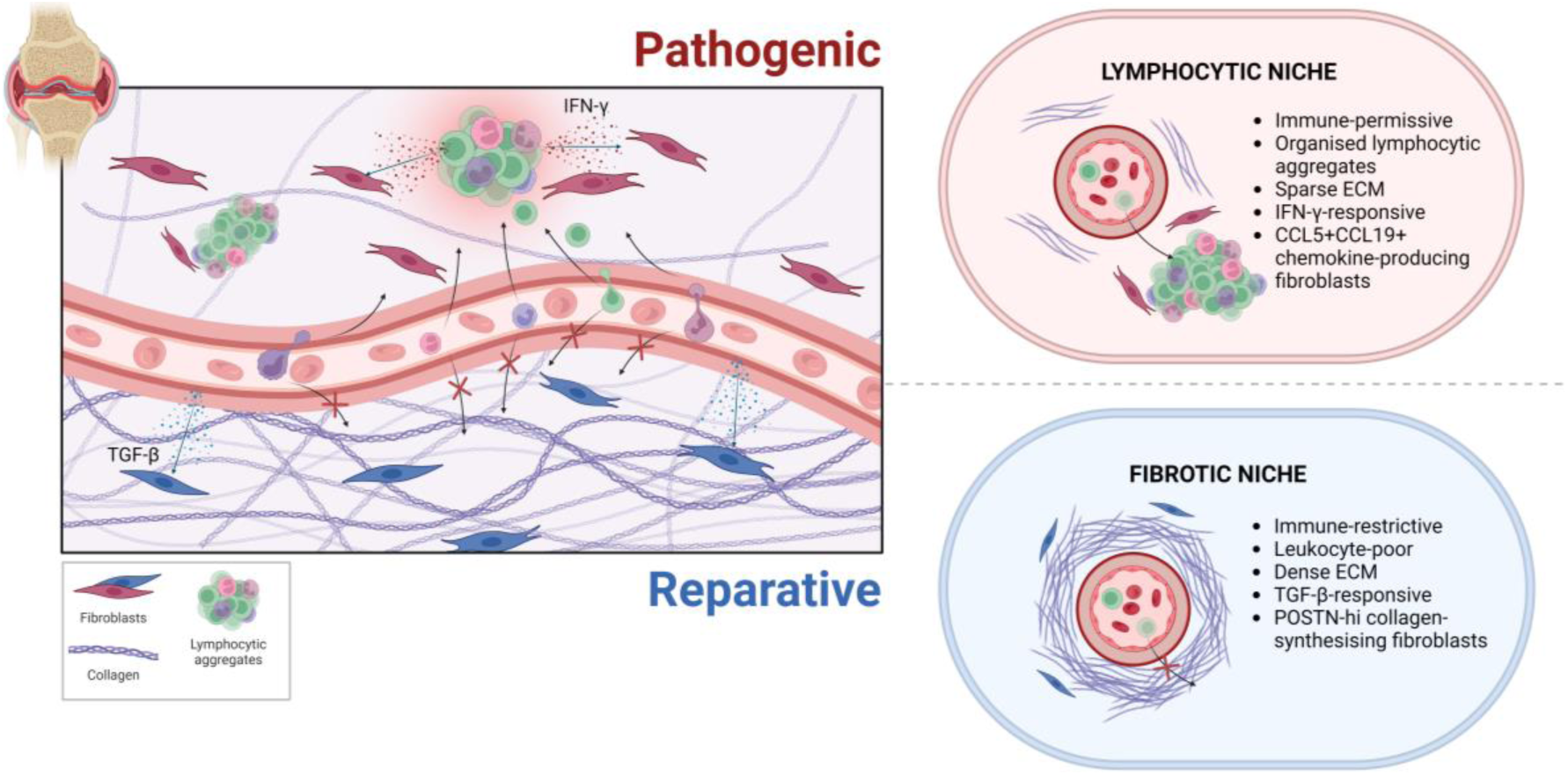

## INTRODUCTION

Immune-mediated inflammatory diseases (IMIDs) are characterised by the persistence of inflammation at anatomically distinct tissue sites^1–3^. Rheumatoid arthritis (RA), a prototypic IMID, is characterised by inflammation localised to the synovial membrane, a thin connective tissue surrounding articular joints^4^. Chronic joint inflammation is associated with the depletion of synovial adipocytes, synovial lining hyperplasia, pathological expansion and activation of sublining fibroblasts, and infiltration of inflammatory cells^5–12^. The cellular and molecular mechanisms underlying this inflammation-driven architectural re-modelling are poorly understood.

Amongst patients with RA, the architecture of the inflamed synovium exhibits heterogeneity in both cellular composition and histological features, with distinct patterns defined as tissue “pathotypes”^13,14^. Pathotype-specific molecular signatures have been identified that demonstrate association with disease outcomes and the likelihood of response to specific therapies^15–19^. Therefore, understanding the disease mechanisms that underpin the development of inflamed tissue architecture may enable development of novel and precision targeted treatments.

Tissue resident fibroblasts are uniquely poised to orchestrate tissue topography by dictating the organisation and composition of immune cells in the tissue. Synovial fibroblasts are phenotypically diverse, their heterogeneity being driven by cell-intrinsic differentiation pathways, tissue-derived molecular signals, and cell-cell communications within the tissue ecosystem^9–17^. While single-cell RNA-sequencing (scRNA-seq) has defined the diversity of the cellular composition of the inflamed synovium, including the heterogeneity of synovial fibroblasts, there is a paucity of data on the spatial distribution of fibroblast states and how location impacts fibroblast-driven pathology. Given the correlation between synovial tissue architecture and disease endotypes, exploring the role of fibroblasts in driving the compartmentalisation of the inflamed tissue microenvironment may enable delineation of the contribution of specific fibroblast phenotypes to inflammatory joint pathology.

Herein, we show that location-defined, cytokine-driven programming of fibroblast identity leads to the construction of distinct tissue niches that regulate the tissue topography. Furthermore, targeted remodelling of the extracellular matrix (ECM) by niche-specific fibroblasts results in either immune-permissive or immune--restrictive tissue niches, dictating immune cell organisation and contributing to the outcome of joint inflammation.

## RESULTS

### The inflamed synovial tissue architecture is defined by the presence of specific cellular niches

To define the functional zonation pattern of the inflamed synovium, we performed probe-based, spatially-resolved transcriptomics (ST) using the 10x Genomics Visium platform. Synovial tissue samples were obtained by minimally invasive, ultrasound-guided biopsies in 4 groups of patients with arthritis, who were naive to treatment with disease modifying anti-rheumatic drugs (DMARDs). These diagnostic groups included: individuals with inflammatory arthritis of ≤3 months symptom duration, who either (1) fulfilled classification criteria for RA by 18 months (early RA) or (2) had an episode of arthritis that spontaneously resolved in the absence of disease activity or DMARD therapy by 18 months (resolving arthritis); (3) individuals with established RA (>3 months’ duration of symptoms at the time of diagnosis and synovial biopsy). (4) Synovial tissue samples were taken from patients with a diagnosis of osteoarthritis (OA) at the time of arthroplasty (**Figure 1a, Supplementary Table ST1**).

**Figure 1.**
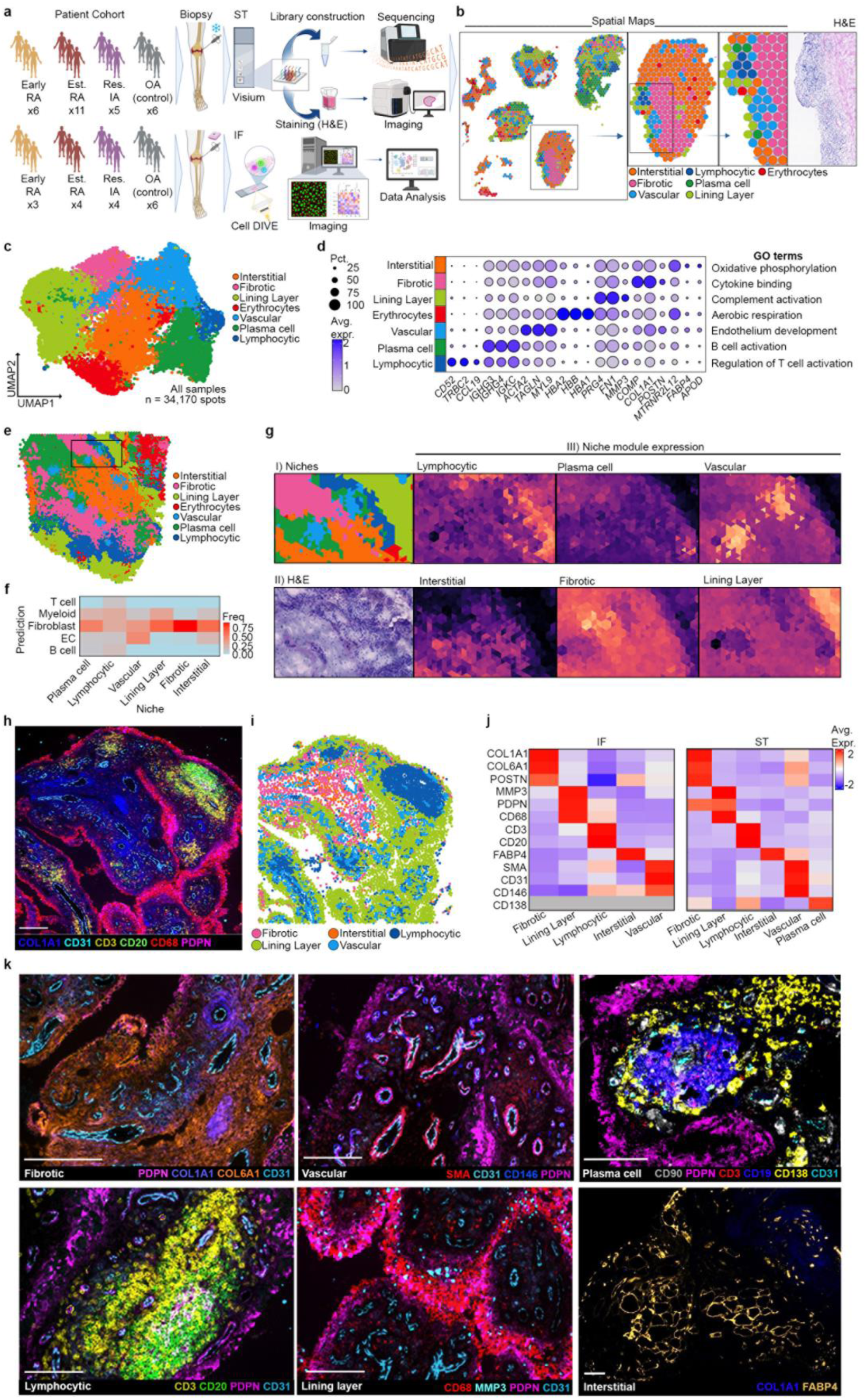
Spatially resolved cellular niches define inflamed synovial tissue architecture. **a)** Schematic overview of multi-modal study design, including patient cohort, sample collection and processing for spatial transcriptomics (ST; 10x Genomics Visium) and multiplex immunofluorescence (IF; Leica Cell DIVE). **b)** Representative spatial plot showing a section of annotated spots from ST of synovial tissue with matched H&E stain. **c)** UMAP representation of ST spots, annotated based on gene expression. **d)** Dotplot of key marker genes and GO terms that define spatial tissue niches. **e)** Spatial plot of deconvoluted spots in synovial tissue following BayesSpace analysis. **f)** Heatmap showing presence of cell types in tisue niches following BayesSpace analysis based on scRNA-seq reference datasets (Zhang et al., 2019; Wei et al., 2020). **g)** Representative spatial plot (I) and matched H&E-stained synovial tissue (II) with spatial plots showing expression of gene modules that define tissue niches (III). **h)** Representative multiplex IF image of synovial tissue showing markers that define tissue architecture. Scale bar: 100 µm. **i)** Matched representative tissue image following computational cell segmentation showing cell centroids and annotation of ST-defined tissue niches. **j)** Comparison of marker gene expression between niches in IF and ST data. **k)** Representative multiplex IF images showing key markers of each niche as well as PDPN, CD68 and CD31 to mark major tissue landmarks. Plasma cell niche imaged using PhenoImager™ HT (Akoya Biosciences); all other niches imaged using Cell DIVE (Leica). Scale bars: 100 µm.

Each tissue section contained 1-8 fragments of biopsied tissue within the ST capture area, taken from distinct locations of an individuals’ synovium to maximise representation of tissue pathology and account for sampling variability (**Figure 1b, S1a, b**). Gene expression of 34,170 capture spots was analysed across 28 patient biopsies (6 early RA, 11 established RA, 5 resolving inflammatory arthritis, and 6 OA patient samples) (**Figure S1a**). Spots were annotated based on their spatial location (**Figure 1b**) and gene expression profiles (**Figure 1c**). Gene Ontology (GO) pathways associated with upregulated genes in these niches further characterised their phenotype (**Figure 1d, S1c**). To determine the cellular composition of these niches, we deconvoluted the gene expression data using 2 publicly available scRNA-seq datasets of human synovial tissue as a reference^812^, to predict the presence of fibroblasts, T cells, myeloid cells, endothelial cells (ECs) and B cells within each ST spot (**Figure 1e-f**).

This combined approach enabled the identification of 6 niches with spatially distinct profiles of gene expression (**Figure 1c-g**): an interstitial niche with high expression of fatty acid transporters and lipoproteins indicative of adipocytes (*FABP4* and *APOD*), enriched in fibroblasts, EC and myeloid gene signatures; a fibrotic niche high in collagen expression (*COL1A1* and *COL3A1*), consisting mainly of fibroblasts; a lining layer niche high in *PRG4* and *FN1* gene expression, enriched in fibroblast and macrophage gene signatures; a vascular niche enriched in ECs, fibroblasts and myeloid cell signatures; a plasma cell niche enriched in immunoglobulin genes (*IGHG3* and *IGHG4*); and a lymphocytic niche with high expression of T cell receptor genes (*TRBC2*) and T and B cell signatures. Erythrocytes, with high haemoglobin gene expression, were also identified and not included in further downstream analysis as these represented small areas of haemorrhage in the tissue from the biopsy.

We confirmed the cellular composition of each niche using multiplex immunofluorescence (IF) imaging (**Figure 1h-j, S2a**) and examined the spatial location of cells within these niches, confirming key architectural and organisational features of the tissue. For example, the tubular structures of vasculature, the continuous barrier of the lining layer cells and aggregation and perivascular cuffing of T / B cells (**Figure 1k**). Collectively, these data demonstrate the presence of distinct synovial tissue niches consisting of specific cellular compositions that define the inflamed synovial architecture.

### Synovial fibroblasts express location-specific gene programs encoded by endothelial cell- and immune cell-derived signals

To determine the identity of fibroblasts within each tissue niche and potential signalling cues responsible for their phenotypes, we used deconvoluted ST gene expression data and performed GO pathway analysis to reveal the distinct transcriptional signatures of fibroblasts within each niche (**Figure 2a, S2b**). Fibroblasts in the fibrotic niche highly expressed ECM-related genes such as *COL1A1* and *COL6A1*; fibroblasts in the lining layer, a well-characterised location-specific synovial tissue niche, highly expressed the genes *PRG4* and *CLU*, in accordance with previous studies^9^; interstitial niche fibroblasts expressed fatty acid-associated genes including *APOD* and *FABP4*; vascular niche fibroblasts highly expressed *NOTCH3* and *ACTA2*, suggesting these fibroblasts likely represent pericytes; and lymphocytic niche fibroblasts highly expressed inflammation-associated genes including *JUNB*, *FOSB* and *FOS* (**Figure 2a**). Fibroblasts in the plasma cell niche could not be adequately deconvoluted due to high expression of immunoglobulin-related genes and were excluded from future analysis of niche-specific fibroblasts.

**Figure 2.**
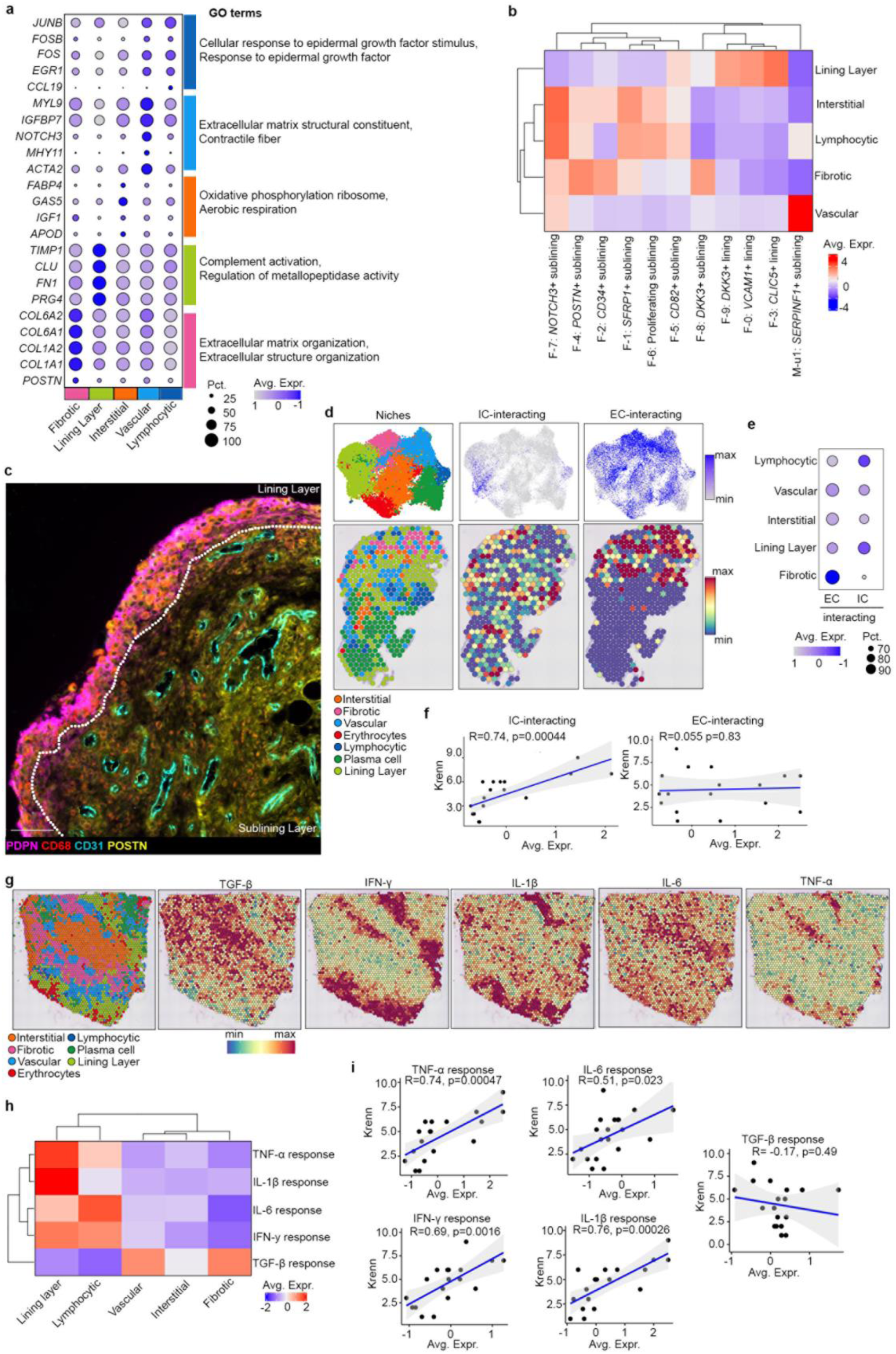
Fibroblast transcriptional identity is defined by spatial microenvironment in synovial tissue. **a)** Dotplot of key marker genes and GO terms in deconvoluted (BayesSpace) fibroblast-rich spots from each niche. **b)** Heatmap showing alignment of fibroblast clusters from scRNAseq analysis of disaggregated synovial tissue (Zhang et al. 2023) to genes associated with fibroblast-rich spots in each niche and a module of the top 40 genes defining the C11 immune cell (IC)-interacting fibroblasts and C4 endothelial cell (EC)-interacting fibroblasts (Korsunsky et al. 2022). **c)** Representative IF image showing perivascular localisation of fibrotic marker POSTN. **d)** Feature plots and spatial gene expression maps demonstrating expression of the immune cell (IC)- and endothelial cell (EC)-interacting gene modules across a synovial tissue section. E) Dotplot showing expression of gene modules of IC-interacting fibroblasts and EC-interacting fibroblasts in fibroblasts from each niche. **f)** Correlation between Krenn histological score and expression of the IC- and EC-interacting gene program in fibroblast-rich spots. Each dot represents a sample. P value calculated using Pearsons’s correlation in ggpubr::ggscatter. **g)** Representative spatial gene expression maps of 5 gene modules defined by genes upregulated in human RA fibroblasts stimulated *in vitro* with indicated cytokines compared to non-stimulated control RA fibroblasts (Tsuchiya et al., 2021). **h)** Heatmap of average scaled expression of cytokine response gene modules in fibroblast-rich spots in each niche. **i)** Correlation between expression of cytokine response gene modules in fibroblast-rich spots and Krenn score. Each dot represents a sample. P calculated using Pearsons’s correlation in ggpubr::ggscatter.

We next mapped niche-specific fibroblast transcriptional identities onto an existing scRNA-seq dataset generated from adult inflamed RA and OA synovium^13^ (**Figure 2b**). We observed that the gene program defining fibroblasts in the fibrotic niche most closely mapped to *POSTN*+, *CD34*+ and *DKK3*+ sublining layer fibroblast subsets (**Figure 2b**). IF staining and RNAscope demonstrated POSTN and *DKK3*, respectively, to be expressed throughout the synovial sublining and enriched in the perivascular space (**Figure 2c, S2c**). In contrast, lymphocytic niche fibroblasts mapped most closely in identity to sublining *NOTCH3*+, *SFRP1*+ and proliferating fibroblasts (**Figure 2c**). NOTCH3-expressing fibroblasts were also located in the perivascular space (**Figure S2d**). Additionally, the vascular niche fibroblasts mapped closely the *SERPINF1*+ mural cell cluster, which further suggested these fibroblasts represent pericytes (**Figure 2c**).

We sought to examine the potential signals responsible for niche-specific fibroblast identities by mapping the previously defined expression pattern of either endothelial cell-interacting (EC-interacting) or immune-cell interacting (IC-interacting) fibroblast genes^20^ in our ST data (**Figure 2d, e**). We observed spatially distinct patterns for each expression program, with fibrotic niche fibroblasts enriched for the EC-interacting program, whilst the lymphocytic and lining layer niche fibroblasts were enriched for an IC-interacting program (**Figure 2d, e**). The IC-interacting signal in the lining layer likely reflects fibroblast-macrophage interactions, where macrophage-specific genes are highly expressed (**Figure S2e**), whereas in the lymphocytic niche this IC-interacting fibroblast program more likely reflects crosstalk between fibroblasts and infiltrating leukocytes in the sublining. Accordingly, in the inflamed synovium reference dataset, *DKK3*+ sublining and *POSTN*+ fibroblasts highly expressed EC-interacting fibroblast genes, whereas *NOTCH3*+ sublining fibroblasts expressed IC-interacting genes (**Figure S2f**).

To validate this, we used matched histology to manually annotate vascular cells, lining layer cells and immune cell aggregates (**Figure S2g**) and examined how the expression of genes associated with niche-specific fibroblasts changed spatially in relation to aggregates and vasculature in ST data. Consistent with IF, distance analysis found that fibrotic niche fibroblast-associated gene expression was highest in the “perivascular” space; meanwhile, lymphocytic niche fibroblast-associated gene expression was highest where aggregates were located and decreased with distance away from aggregates, providing further evidence that fibroblasts respond to polarising signals emanating from these histological features (**Figure S2h**).

Expression of the IC-interacting fibroblast gene program positively correlated with degree of tissue inflammation (measured by Krenn score^21^), unlike expression of the EC-interacting fibroblasts gene program, which demonstrates no such correlation (**Figure 2f**). These data indicate that immune cell infiltration into the tissue triggers a phenotypic shift in fibroblasts, leading them to switch on an immune-interacting phenotype within specific spatial niches in the sublining tissue. This observation suggests a potential feedback loop in which immune cell infiltration actively shapes fibroblast behaviour to contribute to progression of tissue pathology.

### Cytokine priming drives location-specific fibroblast identities within synovial tissue niches

We next sought to determine how interaction with immune cells underpins location-specific fibroblast identities. We examined all upregulated genes (FC ≥2 and adjusted p-value <0.05) from a bulk RNA-seq dataset generated from RA synovial fibroblasts stimulated with various cytokines in culture^22^ to curate fibroblast-specific cytokine response gene signatures and projected these gene signatures onto our ST data (**Figure 2g, h**). We observed spatially distinct patterns of cytokine response by fibroblasts within specific tissue niches. Fibroblasts responding to IFN-γ and IL-6 were expressed in the lymphocytic and lining layer tissue niches, TGF-β-responsive fibroblasts were specific to the fibrotic and vascular niches, while IL-1β and TNF-α responsive fibroblasts were highly expressed in the lining layer (**Figure 2g, h, S3a**).

The cytokine response pattern within spatially defined niches was concordant with that observed in matched fibroblast subsets from scRNA-seq data, with IFN-γ and IL-6 gene response programs enriched in *NOTCH3*+, *CD82*+, *SFRP1*+ and proliferating sublining fibroblasts, whilst lining layer fibroblast subsets were enriched for IL-1β and TNF-α response programs (**Figure S3b**). TGF-β response genes were highly expressed by *POSTN+* and *DKK3+* sublining fibroblasts (**Figure S3b**).

Interestingly, all cytokine response programs in fibroblasts positively correlated with degree of tissue inflammation, except for the TGF-β response program (**Figure 2i**). Therefore, we hypothesised that TGF-β could be a molecular switch that alters fibroblast phenotype towards a fibrotic niche identity which negatively regulates inflammation.

### Niche-specific fibroblasts dictate spatial patterning of immune cell organisation in the tissue through regulation of the matrisome

To examine how TGF-β−responsive fibroblasts in the fibrotic niche might differentially regulate synovial tissue inflammation, we performed an unsupervised analysis of ECM gene expression programs by fibroblasts within each niche, using the Matrisome dataset of 1068 ECM-related genes^23^. We observed increased expression of matrisome genes in the fibrotic niche fibroblasts compared to fibroblasts in all other tissue niches (**Figure 3a**). We next examined the expression of specific matrisome genes across fibroblasts in each tissue niche, revealing distinct programs of ECM-related genes (**Figure 3b, S3c, d)**, and found that fibrotic niche fibroblasts were particularly enriched for genes encoding collagens (**Figure 3c**). These data are consistent with our previous observation that fibrotic niche fibroblasts resemble the *POSTN*+ fibroblast subset, which are TGF-β-responsive and enriched for expression of collagen genes (**Figure S3b, e**).

**Figure 3.**
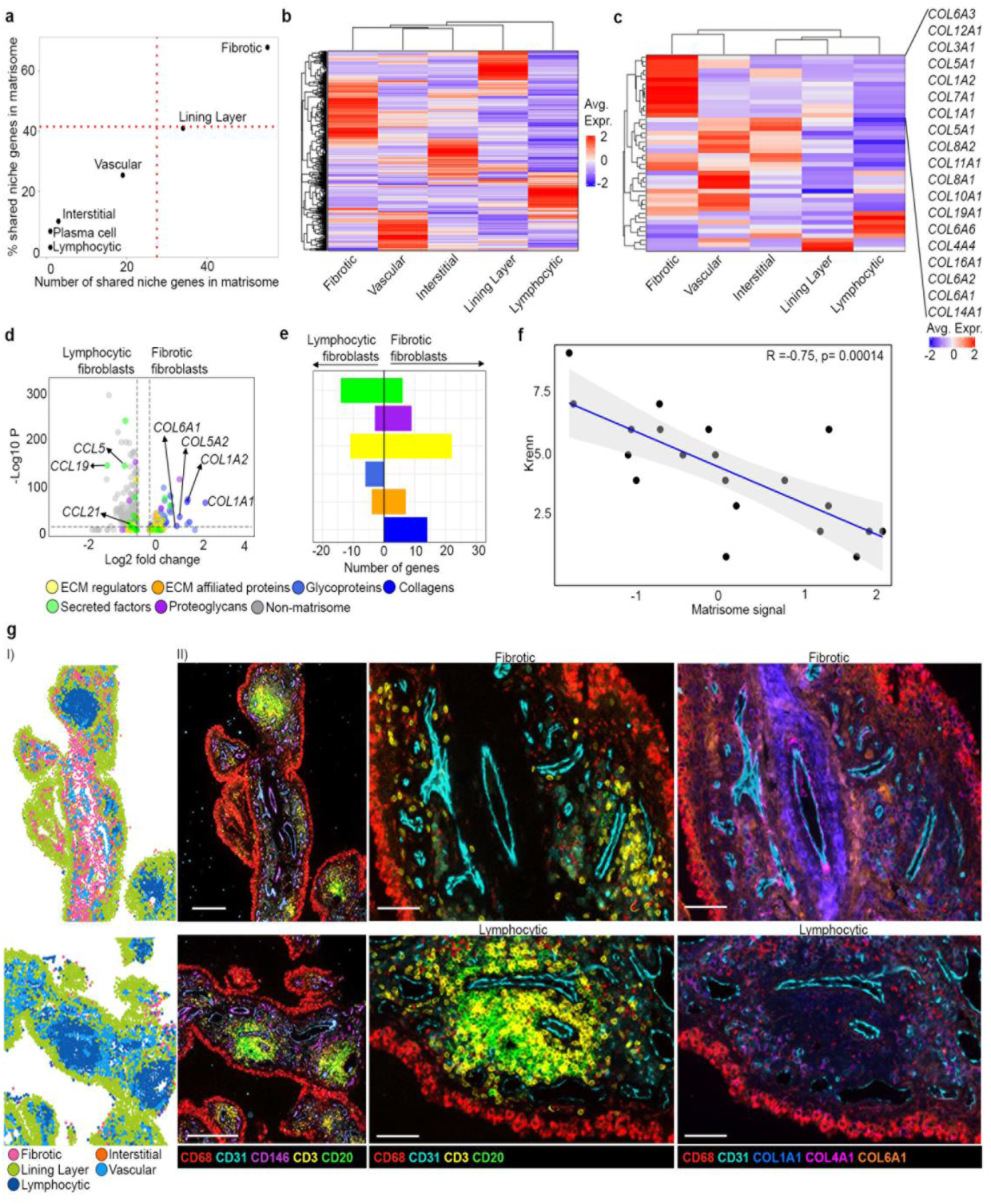
Niche-specific ECM signatures differentially regulate lymphocyte organisation in synovial tissue. **a)** Number of fibroblast genes (x axis) and percentage of fibroblast genes (y axis) in each niche found in the human Matrisome ECM database (Naba et al, 2012; database). **b)** Heatmap of Matrisome ECM genes associated with fibroblast-rich spots in each niche. **c)** Heatmap of collagen genes from the Matrisome enriched in fibroblast-rich spots in each niche. **d)** Volcano plot showing differential expression of ECM-related genes between the lymphocytic and fibrotic niche in fibroblast-rich spots, coloured by Matrisome category. **e)** Number of differentially expressed ECM-related genes between lymphocytic and fibrotic niche in fibroblast-rich spots, coloured by Matrisome category. **f)** Correlation between Matrisome signal, defined by expression of ECM-related genes in ST data, and Krenn score, for each sample. Each dot represents a sample. P value calculated using Pearsons’s correlation in ggpubr::ggscatter. **g)** Multiplex immunofluorescence following cell segmentation showing niche annotations of cell centroids (I) and matched immunofluorescence images (II) (Scale bars: 200µm) showing expression of immune markers or collagen genes in fibrotic and lymphocytic niches, as well as CD68 and CD31 to mark tissue landmarks (II) Scale bar: 50µm.

We then examined whether ECM modulation and collagen deposition underlie distinct tissue niches. We compared differential expression of matrisome genes in fibroblasts from the fibrotic and lymphocytic niches and found an enrichment of collagen genes in the fibrotic niche compared to the lymphocytic niche, alongside an enrichment of chemokines (including *CCL2*, *CCL5*, *CCL19*) involved in leukocyte recruitment and production of ECM-degrading matrix metalloproteinases (MMPs) in the lymphocytic niche (**Figure 3d, e**). Collectively, these data led us to hypothesise that sublining fibroblasts located near vascular endothelium switch on a matrix-synthesising gene expression program in response to TGF-β and deposit specific collagens to construct a fibrotic niche that restricts immune cell trafficking into the tissue. Conversely, in response to IFN-γ and IL-6, immune-interacting fibroblasts present in the lymphocytic niche degrade matrix to facilitate space clearance within the tissue and express leukocyte chemoattractants and survival factors, providing an immune permissive niche that enables aggregation of lymphocytes.

### Collagen deposition within the fibrotic niche forms an immune exclusion zone around blood vessels

We next examined how the construction of a collagen-rich fibrotic niche influences immune cell trafficking into the tissue. Firstly, we observed a negative correlation between degree of synovial inflammation and expression of matrisome genes by fibroblasts (**Figure 3f**), with higher expression of ECM-associated genes by fibroblasts correlating with lower degree of synovial inflammation. We then examined distribution of collagens (collagen I, collagen IV, and collagen VI) that define the fibrotic niche and immune cell aggregates that define the lymphocytic niche using multiplex imaging. We observed formation of collagen-rich barriers surrounding vessels in the fibrotic niche, without nearby immune cell aggregates, whereas vessels in the lymphocytic niche, with nearby immune cell aggregates, had limited expression of collagens I, IV or VI (**Figure 3g**).

Using IF imaging, we confirmed these observations by manually annotating regions around CD31+ vessels and measuring the average staining intensity of CD20, CD3, collagen I, collagen IV, and collagen VI (**Figure 4a**). Collagen I and VI expression was significantly higher in low-immune perivascular regions compared to high-immune perivascular regions (**Figure 4b**). In addition to niche annotation of multiplex IF staining of synovial tissue, we annotated individual cell types based on their expressed markers (**Figure 4c, S4a, b**). Proximity analysis using this annotation confirmed that while COL1-hi, COL6-hi and POSTN-hi fibroblasts are likely found near each other, B/T cell aggregates are not likely to be found near to any of these specific fibroblast populations (**Figure 4d, S4c**).

**Figure 4.**
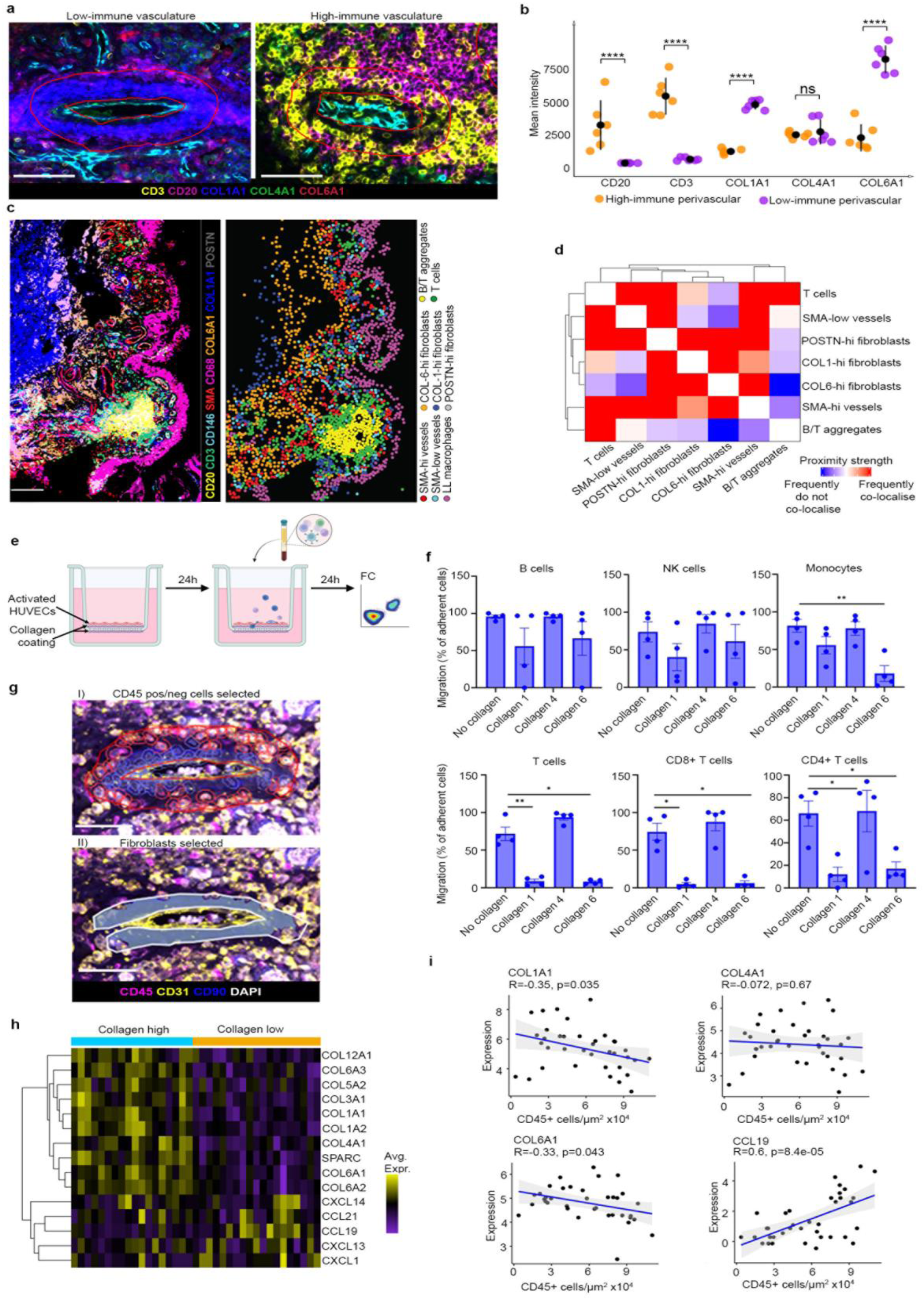
Collagen deposition in the fibrotic niche restricts lymphocyte migration. **a)** Representative immunofluorescence images of areas around vasculature (around n=6 CD31+ vessels from a single sample) with low or high immune infiltration in synovial tissue. Scale bar: 50 µm. **b)** Quantification of immunofluorescence staining intensity of immune and collagen markers. N=6 for high/low-immune perivascular, measured from the same sample. ns, p>0.05; ****, p < 0.0001 (determined by Two-Way ANOVA with Tukey’s HSD Post Hoc tests). Data is mean ± standard deviation. **c)** Annotation of cell types based on marker gene expression in multiplex IF staining (Leica Cell DIVE) of synovial tissue. Scale bar: 100 µm. **d)** Heatmap showing proximity analysis of annotated cell types in multiplex IF. **e)** Schematic overview of transwell migration experiment. **f)** Quantification of transmigrated cells from flow cytometry analysis. N=4 for each condition, across different donors. **, p<0.01; *, p<0.05 (determined by Brown-Forsythe ONE-way ANOVA test with Dunnett’s T3 multiple comparisons test). Data is mean ± SEM. **g)** Representative multiplex immunofluorescence image of synovial tissue analysed using GeoMx® Digital Spatial Profiler (NanoString) showing masks applied to exclude CD45+ cells and select CD90+ fibroblasts to facilitate of collection of transcripts from fibroblasts in regions of interest. Scale bars: 50 µm. Detection of CD45+ cells (red) within 30µm of the endothelial and non-endothelial cells are in circled in grey (I). Detection of CD90+ fibroblasts area selected for RNA probe assessment (greyed area) (II). **h)** Heatmap of expression of indicated genes across regions of interests, showing difference in fibroblast gene expression from areas of high collagen versus low collagen expression. N=18 collagen high regions and n=19 collagen low regions from seven donors. **i)** Correlation between expression of selected genes and number of CD45+ cells x10^4^/µm^2^ within each region of interest of synovial tissue samples. P value calculated using Pearson’s correlation in ggpubr.

To test the hypothesis that specific collagens produced by TGF-β-responsive fibroblasts in the fibrotic niche can limit immune cell migration, we conducted transwell migration assays (**Figure 4e, S5a**) in which ECs were cultured in a monolayer and activated on the apical surface of transwell filters, of which the basal surface was coated with different collagens to mimic vessels surrounded by collagens I, IV and VI. After 24 hours, peripheral blood mononuclear cells (PBMCs) were added to each well and the adherence and migration of T cells, B cells, NK cells and monocytes through the EC- and collagen-lined filters were measured by flow cytometry (**Figure S5a-d**). We observed no change in the proportion of total PBMCs adhering to different collagens compared to control uncoated filters (**Figure S5c**), although a greater proportion of monocytes adhered to the collagen-coated surfaces compared to other cell types. (**Figure S5d**). Compared to wells without a collagen coating, there was a significant decrease in the migration of monocytes (*p* < 0.01), CD4+ and CD8+ T cells (*p* < 0.05) through collagen VI-coated wells, as well as significantly decreased CD4+ and CD8+ T cell migration through collagen I-coated wells (*p* < 0.05) (**Figure 4f**). These data suggest the presence of collagens I and VI restrict migration of specific leukocytes through ECs and into the tissue.

To validate the role of perivascular collagens in restricting leukocyte infiltration, we undertook targeted spatial transcriptomics (Nanostring GeoMX® Digital Spatial Profiler) of human synovial tissue to sequence CD90+ fibroblasts from perivascular regions with varying levels of CD45+ cell infiltration (**Figure 4g**). Global analysis revealed collagen high and collagen low perivascular regions, again demonstrating a distinction between fibroblast populations enriched for either collagen expression or production of chemokines (**Figure 4h**). In line with the results from our transwell assay, we found that perivascular fibroblast expression of *COL6A1* and *COL1A1* negatively correlated with CD45+ cell number per area, although no such correlation was observed with *COL4A1* expression (**Figure 4i**). As expected, *CCL19* expression positively correlated with number of CD45+ cells (**Figure 4i**). These data confirm that collagens, notably collagen I and VI, deposited by fibrotic niche fibroblasts are associated with restriction of immune cell migration into the synovial tissue.

### Endothelial cell-derived TGF-β production via NOTCH signalling drives collagen deposition and fibrotic niche fibroblast identity

We next investigated the ability of ECs to produce TGF-β to which fibrotic niche fibroblasts respond. Distance analysis demonstrated that the TGF-β response module was most highly expressed in fibroblasts in close proximity to blood vessels and decreased in signal intensity towards the lining layer (**Figure 5a**). Therefore, we analysed publicly available gene expression data from synovial organoids, in which RA synovial fibroblasts were cultured either alone or with ECs^12^ (**Figure 5b**). The analysis revealed that co-culturing ECs with synovial fibroblasts in organoids led to increased expression of TGF-β response genes, fibrotic niche fibroblast genes, and expression of collagens (*COL1A1*, *COL4A1* and *COL6A1*) (**Figure 5b, c**). Accordingly, IF staining of organoids demonstrated significantly increased expression of collagen I, collagen VI and POSTN in close proximity to endothelial cells in fibroblast-EC organoids compared to fibroblast-only organoids (**Figure 5d**). Treatment of fibroblast-EC organoids with DAPT, a NOTCH inhibitor, demonstrated that induction of the fibrotic niche fibroblast phenotype was NOTCH-dependent (**Figure 5b, c**). Furthermore, analysis of our Nanostring GeoMx data demonstrated that expression of genes upregulated by NOTCH signalling negatively correlated with number of CD45+ cells (**Figure S4d**). Taken together, these data suggest that EC-driven priming of fibroblasts via NOTCH signalling is sufficient to induce fibrotic niche identity and expression of collagens in perivascular fibroblasts at the transcript and protein level, even in the absence of immune cells. Moreover, in tissue, expression of NOTCH-dependent genes correlates with restriction of immune infiltration.

**Figure 5.**
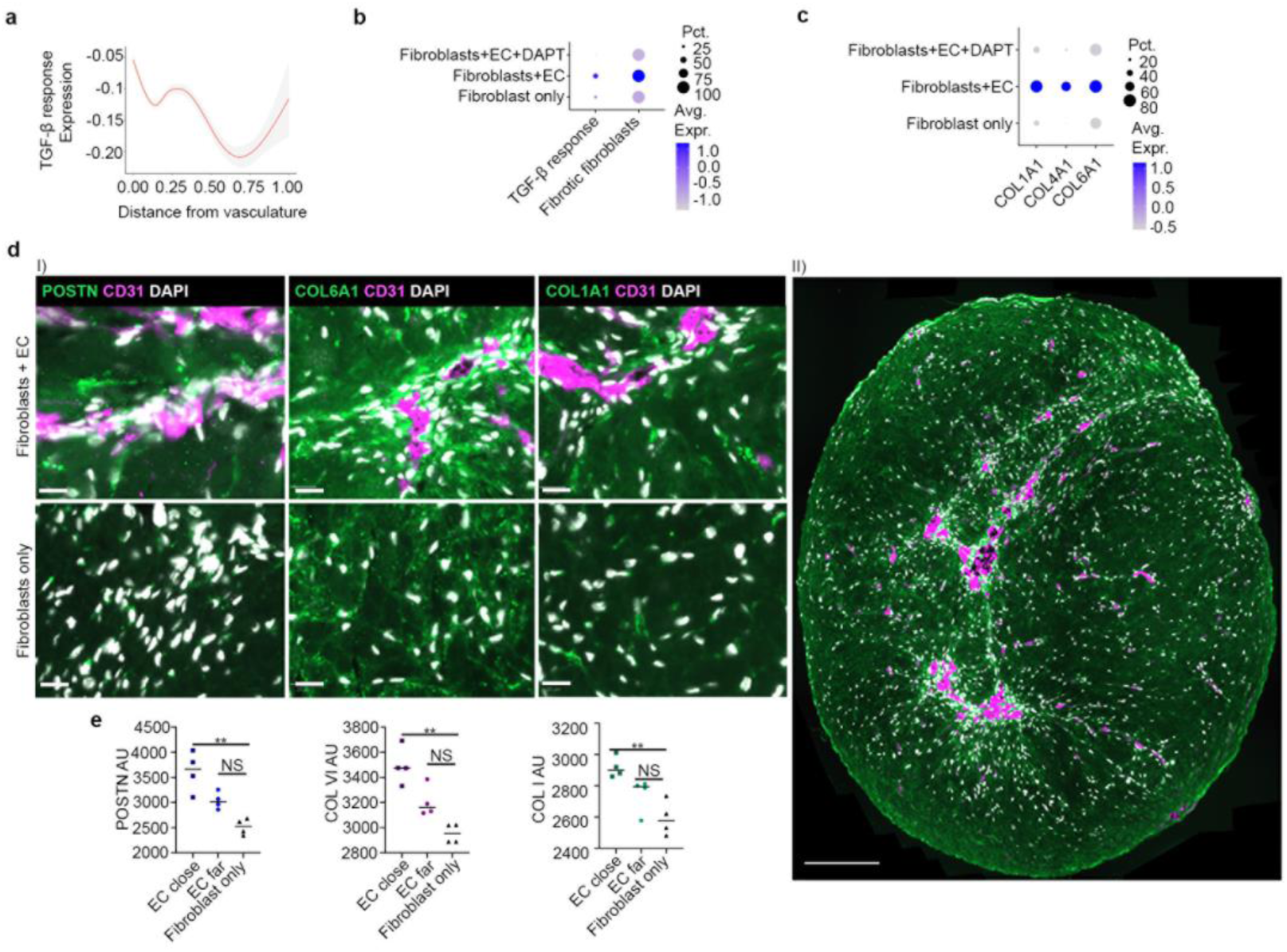
Synovial organoids demonstrate EC-derived TGF-β signalling via NOTCH. **a)** Distance analysis showing expression of TGF-β fibroblast response module genes from vasculature towards the lining layer. **b)** Dotplot showing expression of TGF-β response module and fibrotic fibroblast marker genes in synovial organoid data from Wei et al., 2020. **c)** Dotplot showing expression of selected collagens in organoid data from Wei et al., 2020 (Fibroblast only: organoid containing fibroblasts only; Fibroblasts+EC: organoids containing fibroblasts and ECs; Fibroblasts+EC+DAPT: organoids containing fibroblasts and ECs treated with a NOTCH inhibitor). **d)** Immunofluorescence images (I) Scale bars: 20µm, (II) Scale bars: 200µm and **e)** quantification of POSTN, Col VI or Col I staining in organoids containing fibroblasts only or fibroblasts and ECs, either “close” or “far” in respect to ECs. N=3 for each condition. **, p<0.001; NS, p>0.05, as determined by Kruskal-Wallis test with Dunn’s post-hoc test. EC, endothelial cell.

### A common POSTN+ ECM-remodelling fibroblast gene program is implicated in negative regulation of inflammation in IMIDs and cancer

Given the differential effect of niche-specific fibroblasts on immune cell organisation within the tissue, we postulated that these fibroblasts could influence the outcome of tissue inflammation. Firstly, we investigated the involvement of the fibroblasts in our ST niches across different stages of RA by analysing scRNA-seq data from individuals with active RA, those in remission and healthy donors^24,25^ (**Figure 6a, S7a**). We observed increased expression of the fibrotic fibroblasts, EC-interacting and TGF-β response genes in the THY1^high^ sublining fibroblasts, which have been shown to be associated with expression of collagens and ECM-related genes^24^ (**Figure 6a**). Interestingly, these gene programs were not expressed in healthy samples but highly expressed in active RA, with some expression in remission, suggesting this fibroblast phenotype is induced in response to inflammation and may contribute to an active repair process in remission. In contrast, the interstitial niche fibroblast genes were more highly expressed in healthy synovium, likely due to the presence of adipocytes, which is consistent with previous molecular mapping studies of healthy synovium^5^.

**Figure 6.**
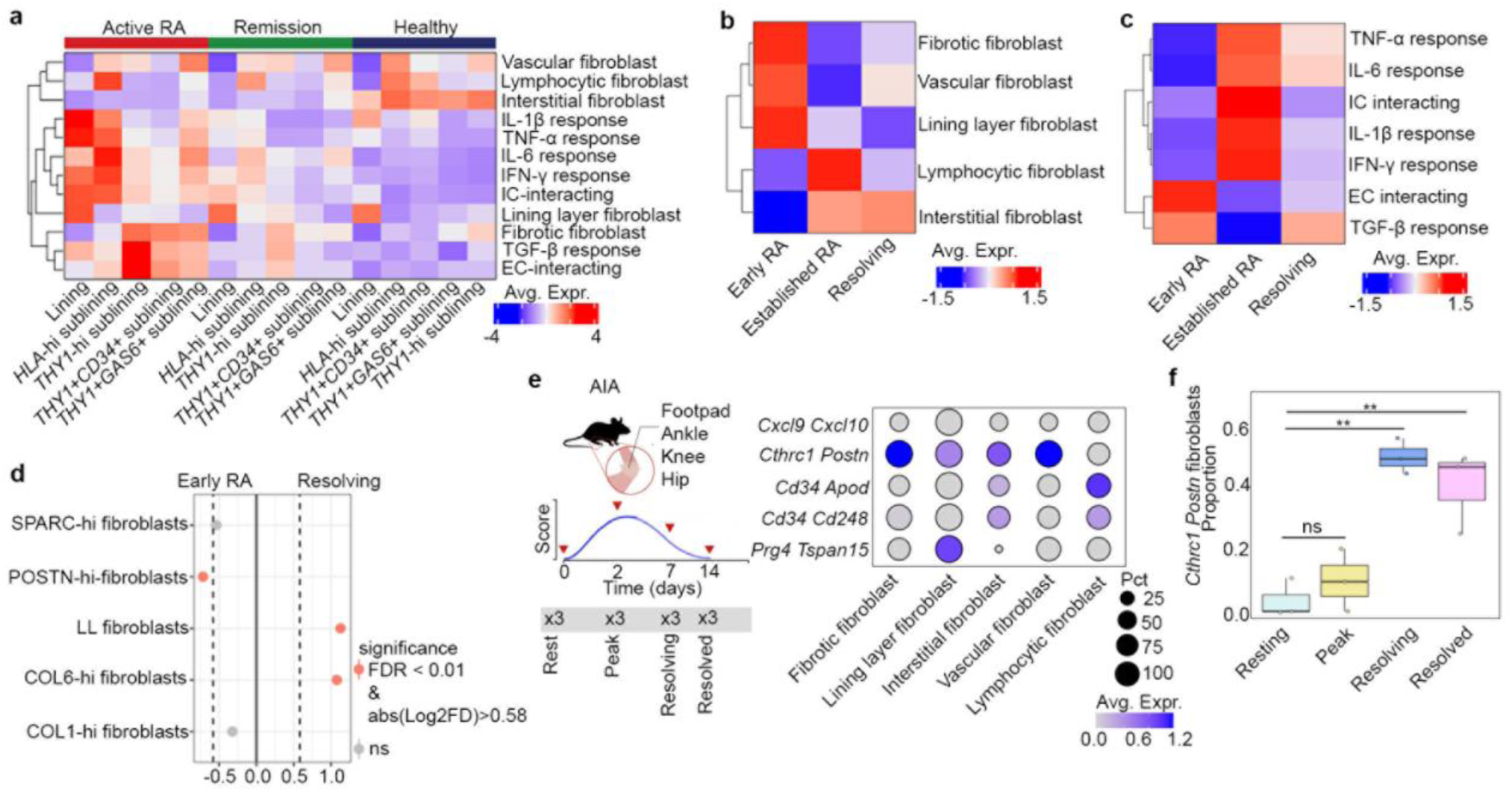
ECM-remodelling fibroblasts are induced in response to inflammation to promote tissue repair and modulate immune restriction. **a)** Heatmap showing expression of niche-specific fibroblast genes, cytokine response and IC- and EC-interaction programs across fibroblast clusters from healthy, active RA and remission synovial tissue samples (Alivernini et al. 2020). **b)** Heatmap of expression of genes from fibroblast-rich spots in each niche in early RA, established RA and resolving arthritis samples. **c)** Heatmap of fibroblast cytokine response and EC- and IC-interaction programs in early RA, established RA and resolving arthritis samples. **d)** Relative niche abundance of annotated cell types from multiplex IF between early RA and resolving arthritis samples. **e)** Overview of antigen-induced arthritis (AIA) timecourse data and expression of niche-specific fibroblasts gene programs in AIA fibroblast clusters. **f)** Relative proportion of Cthrc1+ Postn+ cluster across the AIA timecourse. NS, p>0.05. **, p<0.01. Significance determined by Brown-Forsythe ONE-way ANOVA test with Dunnett’s T3 multiple comparisons test. Box plot represents median with lower and upper hinges correspond to the first and third quartiles. The upper/lower whisker extends from the hinge to the smallest/largest value no further than 1.5 * IQR from the hinge.

Although comparison of niche abundance in early RA, resolving arthritis, established RA and OA revealed few differences at the spatial transcriptomic level, IF analysis revealed that, compared to early RA, established RA had a higher abundance of the lymphocytic and lining layer niches, while early RA had a higher abundance of the interstitial niche, consistent with the depletion of adipocytes, infiltration of immune cells and synovial lining hyperplasia that are key architectural features of inflamed synovial tissue (**Figure S6a, b**). Interestingly, the fibrotic niche showed increased abundance in early RA compared to either resolving arthritis or established RA, as well as increased abundance in OA compared to established RA (**Figure S6b**).

We next evaluated niche-specific fibroblast gene expression across disease groups. Concordant with our previous findings, the lymphocytic niche fibroblast genes were most highly expressed in established RA, suggesting a role for lymphocytic fibroblasts in sustaining chronic inflammation, while fibrotic niche fibroblast genes were most highly expressed in early RA (**Figure 6b**).

We also investigated cytokine response programs in fibroblasts across disease groups and found, as expected, increased expression of all cytokine response gene programs in established RA, except for TGF-β which was increased in early RA and resolving arthritis (**Figure 6c**). Similarly, expression of the IC-interacting gene program was more associated with established RA, whereas the EC-interacting program was most associated with early RA (**Figure 6c**).

We also performed analysis stratified by RA synovial tissue pathotype, focussing on the differences between tissues defined as lymphoid (defined by presence of ≥1 small aggregate in 2 tissue fragments or at least 1 larger aggregate, determined by aggregate radial size; n = 10) and pauci-immune (defined by not reaching lymphoid criteria and low infiltrate density score; n = 7). We found that the key differences between the fibrotic and lymphocytic niche were also observed at the global tissue level between these two pathotypes (**Figure S6c-f**). Lymphoid samples demonstrated increased expression of the IC-interacting fibroblast gene program, while pauci-immune samples highly expressed the EC-interacting fibroblast gene program (**Figure S6c**). Lymphoid samples were associated with increased expression of IFN-y, IL-1β, TNF-α and IL-6 response programs, whereas the pauci-immune samples were associated with TGF-β response (**Figure S6d**). Differential gene expression analysis of matrisome genes showed that fibroblasts in pauci-immune tissues had significantly increased expression of *POSTN* and *COL6A1*, while fibroblasts in lymphoid tissues had significantly increased expression of *MMP1* and *MMP3*, alongside chemokine gene expression (*CCL19* and *CXCL1;* **Figure S6e, f**). Analysis of niche abundance showed a modest increase in the proportion of the lymphocytic niche in lymphoid samples compared to pauci-immune samples, and, accordingly, niche-specific fibroblast abundance analysis demonstrated that lymphoid tissues were associated with lymphocytic and lining layer niche fibroblast gene expression, while pauci-immune tissues were associated with fibrotic, interstitial and vascular fibroblast gene expression (**Figure S6g-h**).

To investigate how differences in synovial tissue composition determines disease trajectory, we compared abundance of cell types visualised by multiplex IF between early RA and resolving arthritis (**Figure 6d**). We found that POSTN-hi fibroblasts were enriched in early RA, while collagen VI-hi fibroblasts were enriched in resolving arthritis, implying a role for collagen VI-producing fibroblasts in the successful resolution of inflammation (**Figure 6d**). This is consistent with our observation that collagen VI deposition underpins a pro-resolution process in the joint.

To enable temporal analysis of niche fibroblast abundance during the active resolution phase of arthritis, we next analysed our niche-specific fibroblast signatures in a scRNA-seq dataset of mouse antigen-induced arthritis (AIA) (**Figure 6e, f, S7b**). We observed that the fibrotic niche fibroblast signature was most highly expressed in *Cthrc1*+ *Postn*+ fibroblasts (**Figure 6e**) which were significantly expanded in the resolving and resolved phases of arthritis compared to the resting state (**Figure 6f**).

Comparing the phenotype of the fibrotic niche fibroblasts to a cross-tissue fibroblast atlas^20^ showed they corresponded to vascular-interacting fibroblast phenotypes (*CD34*+*MFAP5*+ C9, *SPARC*+*COL3A1*+ C4) common across several tissues (**Figure S7c**). Interestingly, gene expression analysis of cancer-associated fibroblasts in breast cancer^26^ demonstrated that the fibrotic niche fibroblast gene expression program was more expressed in myofibroblast cancer-associated fibroblasts (myCAF) than inflammatory cancer-associated fibroblasts (iCAF) (**Figure S7d**). Moreover, this signature also mapped to the *LRRC15*+ myofibroblast cluster found in multiple human cancers, which initiates TGF-β-dependent ECM-remodelling and dampens anti-tumour immunity through suppression of CD8+ T cells^27,28^ (**Figure S7e**). Furthermore, examining expression of canonical myofibroblast-related genes, including *MFAP4*, *TNC*, *FMOD*, *CTHRC1*, *POSTN* and *LRRC15*, across our niche-specific fibroblasts showed the fibrotic niche fibroblasts are associated with expression of these genes (**Figure S7f**).

Together, these data suggest that deposition of collagens by TGF-β-responsive POSTN+ fibroblasts represent a common myofibroblast-like state, modulating an immune-restricting tissue process that operates in inflamed tissue to resolve inflammation.

## DISCUSSION

Our previous work has uncovered significant fibroblast heterogeneity, including the existence of pathogenic synovial fibroblasts with non-overlapping effector functions responsible for either inflammation or damage that reside in distinct anatomical compartments of the synovium^9^. More recently, regulatory fibroblast populations, which promote resolution of inflammation through induction of pro-resolving immune cell populations have been identified^29^. Herein, we describe fibroblast heterogeneity in the context of spatially relevant tissue topography, identifying a novel mechanism of fibroblast-mediated resolution of inflammation via ECM remodelling to create immune exclusion zones in the tissue.

We describe, near lymphocytic aggregates, fibroblasts that are capable of amplifying inflammation through upregulation of genes associated with T cell recruitment and survival as well MMP production (*CCL5*, *CCL19* and *CCL21*). Our analysis suggests IFN-γ and IL-6 derived from infiltrating leukocytes drives a molecular switch in these lymphocytic niche fibroblasts towards an immune-interacting phenotype. In secondary lymphoid organs, specialised fibroblasts orchestrate lymphocyte compartmentalisation, producing distinct combinations of chemokines and depositing complex ECM that serves as a scaffold along which dendritic cells and lymphocytes engage and migrate to facilitate adaptive immune responses^30^. Similarly, we propose that the lymphocytic niche fibroblasts we identified underpin persistence of joint inflammation by facilitating recruitment and aggregation of lymphocytes in the tissue, evidenced by a strong correlation of abundance of IFN-γ- and IL-6-responsive lymphocytic niche fibroblasts with severity of joint inflammation, duration of disease, and association with lymphoid synovial tissue pathotype.

IFN-γ-responsive fibroblasts have been reported to be a common pathogenic fibroblast state in the inflamed target tissues of multiple IMIDs^20^. HLA-DR+ sub-lining fibroblasts were be expanded 15-fold in RA synovium and represent an activated fibroblast state induced by IFN-γ signalling^8,31^. *In vitro* studies have shown that HLA-DR+ fibroblasts are capable of presenting antigens to CD4+ T cells^32–34^ and secrete pro-inflammatory chemokines and cytokines that drive a positive feedback mechanism, promoting recruitment of diverse immune cell types into the inflamed tissue^8,13^. A transition from an immunosuppressive to an intrinsically immune-stimulatory phenotype in early arthritis has been previously observed *in vitro* during the first three months of treatment^35^. We, therefore, propose that the subtle infiltration of T cells into the synovium observed in pre-clinical phases of RA^36^ represents an early disease mechanism, that, when combined with successful fibroblast activation via IFN-γ, results in the construction of an immune-permissive niche, leading to a feedforward mechanism that amplifies lymphocyte recruitment to promote chronic joint inflammation.

In contrast, we identified a fibrotic tissue niche associated with a paucity of immune cells and defined by the presence of TGF-β−responsive, ECM-remodelling fibroblasts that deposit a collagen-rich barrier in close proximity to blood vessels. These TGF-β−responsive fibroblasts mapped to previously identified *POSTN*+ and *DKK3*+ synovial fibroblast phenotypes that have been shown to be enriched in ECM gene expression programs and tissue remodelling pathways^13,37^. Consistent with these observations, we found that TGF-β−responsive fibroblasts in the fibrotic niche express a specific collagen gene expression profile, like myofibroblasts in other tissue sites and myCAFs in tumours^26,37^. The collagen-rich barrier contains collagens I and VI that were each able to significantly inhibit trans-endothelial leukocyte migration *in vitro*. A similar mechanism has been shown to contribute to immune cell exclusion in tumours, driven by myCAFs, which have been recognised as pivotal contributors to the blockade of T cell infiltration via deposition of highly organised ECM^38–40^. We conclude, therefore, that collagen-rich barriers around blood vessels deposited by TGF-β−responsive fibroblasts represented an immune exclusion zone constructed as part of a reparative process to limit further lymphocyte migration into the tissue.

Our analysis shows that the ECM-remodelling TGF-β−responsive fibroblast state was only observed in inflamed and not in healthy synovial tissue and were only significantly expanded during the resolution phases of experimental arthritis in mice. This suggests that these fibroblasts represent an emergent state, activated in response to inflammation, which act in a negative feedback loop to restrain inflammation. Our data found that this fibroblast state was regulated by TGF-β signalling in a NOTCH-dependent manner, consistent with their proximity to vasculature, as vascular endothelium-derived NOTCH ligands are known to regulate positional identity of perivascular fibroblasts in the synovial sublining^12^. Although we observe here that signalling from endothelial cells was sufficient to drive the fibrotic niche fibroblast state, it is likely that the source of TGF-β in the tissue is multi-cellular, and the mechanisms governing TGF-β signalling in response to inflammation in the synovium require further investigation.

An ECM-remodelling TGF-β−responsive myofibroblast-like state has been reported in studies of other tissues, including recent lineage tracing studies in experimental lung fibrosis models, where TGF-β signalling promoted the emergence of a Cthrc1+ Postn+ fibroblast state following injury, which mediated tissue fibrosis and suppressed the emergence of an inflammatory fibroblast state^37^. Crucially, blocking the induction of this fibrotic phenotype exacerbated inflammation^41^. Furthermore, in cancer, TGF-β signalling is critical for the emergence of cancer-associated LRRC15+ myofibroblasts that suppress CD8+ T cell responses^28^. While lineage tracing studies are required to confirm this in the synovium, collectively, these data support the paradigm of multiple potential fibroblast differentiation trajectories that emerge under the direction of specific microenvironmental signalling cues, including a common TGF-β-driven myofibroblast-like state that restrains inflammation.

Our data suggests that the construction of distinct tissue niches in the synovial tissue may be a key determinant of tissue pathotype, with important implications for disease outcomes and therapy response. The pauci-immune synovial tissue pathotype is associated with robust expression of a canonical fibroblast transcriptional signature and scant infiltration of immune cells^14,42,43^. Importantly, RA patients with this pathotype are less likely to respond to DMARDs or biologic therapies^42^. We demonstrated that pauci-immune synovial tissue is associated with expression of TGF-β-responsive ECM-remodelling fibroblast genes, and scant leukocyte infiltration in this tissue is consistent with our observation of the immune-restrictive role of these fibroblasts. The pauci-immune tissue pathotype may, therefore, represent synovial tissue in which fibrotic niche fibroblasts successfully mediate immune exclusion, but symptoms (notably fatigue and pain) driven by other disease mechanisms persist, underpinning pathology in these patients^42,44^. Further investigation into the non-inflammatory processes promoting disease in patients with pauci-immune synovial tissue may highlight novel therapeutically targetable pathways to address the unmet clinical need in these patients.

In summary, we propose that perivascular fibroblasts in the synovium are poised to differentially respond to signalling cues from the vasculature or infiltrating immune cells, leading to the emergence of distinct fibroblast states that construct sub-synovial tissue niches which are either restrictive or permissive to immune cell infiltration to influence the outcome of joint inflammation. Furthermore, these data suggest that fibroblasts undertake collagen synthesis in response to inflammation as an attempt to “heal”, leading to immune cell exclusion and limited fibrotic scar formation. Development of treatments that augment endogenous fibroblast-driven tissue repair and regulatory mechanisms to re-establish homeostasis under inflammatory conditions may offer novel therapeutic approaches to restoring diseased tissue microenvironments to a healthy state.

## METHODS

### Patient Demographics and Ethical Approval

Synovial tissue samples were obtained by minimally invasive ultrasound-guided synovial tissue biopsies from patients in the Birmingham Early Arthritis Cohort (BEACON) study^45,46^. Patients with clinical synovitis were recruited to the BEACON cohort upon rheumatologists’ assessment of symptoms attributed to inflammatory joint disease (pain, stiffness and/or swelling in at least one synovial joint). Samples included in this study were defined as early RA (≤3 months symptom duration), established RA (>3 months symptom duration) or resolving inflammatory arthritis (no clinical evidence of synovial swelling at follow-up assessment and had not taken DMARDs or received glucocorticoid treatment in any form in the preceding 3 months prior to final follow-up). Patients were classified as having RA according to the 1987 and/or 2010 ACR/EULAR classification criteria for RA. Samples from osteoarthritis (OA) patients were obtained from arthroplasty procedures. All patients were naïve to treatment with disease-modifying antirheumatic drugs (DMARDs) at inclusion.

The BEACON study (12/WM/0258) and the study recruiting patients undergoing arthroplasty for osteoarthritis (07/H1204/191) were approved by the West Midlands Black Country research ethics committee. All patients gave written, informed consent.

### Histological Assessment of Synovial Tissue

Haematoxylin and Eosin (H&E) stained synovial biopsy sections were examined histologically for the severity of inflammatory infiltrate using the inflammatory component of the Krenn synovitis score^21^. Inflammatory infiltrates were graded from 0 to 3 (0 = no inflammatory infiltrate, 1 = few mostly perivascular-situated lymphocytes or plasma cells, 2 = numerous lymphocytes or plasma cells sometimes forming follicle-like aggregates, and 3 = dense band-like inflammatory infiltrate or numerous large follicle-like aggregates).

To classify the samples into pathotypes on the basis of H&E staining, semi-quantitative four-point scores for infiltrate density and aggregate radial size were used^39^, as validated by scoring tissues from the phase 1 AMP RA cohort^8^ and implemented in the AMP2 cohort^13^. Aggregate grade was derived as follows: Grade 3; high ≥ 20 radial count. Grade 2: medium 10–19 radial count. Grade 1: low 6–9 radial cell count. Grade 0: No aggregates. Global density of infiltrating cells was graded using an atlas on a semiquantitative 0-3 scale. This grading system was used to classify the samples into three pathotypes - either lymphoid, diffuse or pauci-immune according to the following rules. Lymphoid: The presence of ≥1 grade 1 aggregate in at least two fragments, or any grade 2 aggregate, or any grade 3 aggregate. Diffuse: Does not meet lymphoid criteria but with a mean fragment density score ≥ 1. Pauci-immune: Does not meet lymphoid criteria, mean fragment infiltrate density score < 1.

### Synovial Tissue Processing and Sample Preparation for Visium Spatial Gene Expression (10x Genomics)

Synovial tissue samples were frozen in Tissue-Tek OCT Medium (4583, Sakura) after collection. Cryostat sections were cut at 10μm onto Visium Spatial Gene Expression Slides (PN-1000185, 10x Genomics) to fit the 4 6.5 x 6.5mm spatial transcriptomic capture areas. Sections were incubated for 1 minute at 37 °C and immersed in pre-chilled (−20 °C) methanol for fixation for 30 minutes, and then stained with H&E, according to 10x Genomics protocols. Each capture area was imaged using Axio Z1 Slide Scanner (Zeiss).

### Library Construction and Sequencing for Visium Spatial Gene Expression (10x Genomics)

Permeabilisation of the synovial tissue samples was performed for a total of 12 minutes at 37 °C, and libraries were constructed using the Visium Spatial Gene Expression Library Construction Kit (PN-1000186 and PN-1000190, 10x Genomics), according to manufacturers’ instructions. Library quantification was carried out using KAPA SYBR FAST kit (KK4600, Sigma-Aldrich) and sequenced by Genomics Birmingham (Birmingham, UK) using a high output reagent Dual Index Kit TT Set A (10x Genomics) and the NextSeq500 machine (Illumina).

### Computational Analysis of Visium Spatial Gene Expression (10x Genomics)

Sequenced reads were aligned to GRCh38, and count matrices generated using Space Ranger (10X Genomics, v1.3.0) and read into RStudio (v4.3.1). Data was processed using Seurat (v4.3.1) and sctransform (v0.3.5). Batch correction was applied using Harmony (v0.1.1) and niches were defined using Seurat::FindNeighbours() and Seurat::FindClusters(). A cluster resolution of 0.2 was selected and marker genes identified using Seurat::FindAllMarkers(). BayesSpace (v1.6.0) deconvolution was also performed. Distances from spatial landmarks were analysed using Sp (v2.0-0). GO term enrichment analysis was completed using gsfisher (v0.2).

### scRNA-seq Analysis

The following quality control (QC) metrics were used for data from Tang et al., 2024: nFeature_RNA > 200 & nFeature_RNA < 6000 & percent.mt < 10. Data was processed using the following Seurat (v4.3.1) functions: NormalizeData(), FindVaribleFeatures(), ScaleData(), RunPCA(), RunUMAP(), FindNeighbours(), FindClusters(). Samples were integrated using FindIntegrationAnchors() and IntegrateData(). Fibroblasts from Tang et al., 2024, identified on the basis of gene expression, were labelled with fibroblast cluster identities from Alivernini et al., 2020 using FindTransferAnchors(). Data were harmonised (where required) using Harmony (v0.1). Volcano plots were plotted using EnhancedVolcano (v1.13.2) and heat maps were generated using ComplexHeatmap (v2.10) R packages.

### Manual annotation of architectural landmarks in synovial tissue for ST analysis

Vascular, lining layer and aggregates were manually annotated by referencing matched H&E-stained synovial tissue. Perivascular zones were defined as spots within 5% of the total distance from vascular manual annotations, and the peri-vascular zone defined as the spots within 5-15% of the total distance from vascular manual annotations. The immune cell aggregates zone was defined as spots within 5% of the total distance from aggregate manual annotations and the peri-aggregate zone was defined as the spots within 5-15% of the total distance from aggregate manual annotations.

### Synovial Tissue Processing for Multiplex Immunofluorescence Staining (Leica Cell DIVE and Akoya Biosciences PhenoImager^TM^)

Matched formalin-fixed paraffin-embedded biopsy tissue were used for proteomic validation of ST findings. For Cell DIVE (Leica), paraffin-embedded synovial tissue was sectioned at 4 μm, incubated for 1 hour at 60°C, deparaffinised and rehydrated by following the Cell DIVE standardised protocol consisting of successive changes of Xylene (2x wash) for 5 minutes each with gentle agitation, followed by 2x wash of 5 minutes each in 100%, 95%, 70% and 50% ethanol all with gentle agitation, and two changes of PBS for 5 minutes each. Permeabilisation was performed for 10 minutes in 1X PBS with 0.3% Triton X100, followed by a PBS wash of 5 minutes. Antigen retrieval was carried out using a pressure cooker, according to manufacturer’s recommendations. Slides were then stained with DAPI and imaged for the first scan plan. Imaging using Cell DIVE was performed at 20X to acquire background autofluorescence and generate virtual H&E images. Each Cell DIVE staining round consisted of 3 conjugated antibodies incubated at 4°C overnight or for an hour at room temperature. Manually conjugated antibodies were purchased in a BSA-Azide-free format and conjugated using antibody labelling kits (A20181, A20187, A20186, ThermoFisher). Full antibody details can be found in **Supplementary Table ST12**. Between staining rounds, slides were bleached and re-stained with DAPI to assist in image registration and alignment.

For assessment of plasma cells, the PhenoImager™ HT (Akoya Biosciences) system at Birmingham Tissue Analytics was used. Sections were stained using the BOND RX Autostainer (Leica) platform using sequential staining. Dewaxing was performed using BOND Dewax solution (AR9222, Leica) and antigen retrieval was performed at both pH9 and pH6. To block non-specific staining, 1X Antibody Diluent/Block solution (ARD1001EA, Akoya Biosciences) was used. Primary antibodies were detected using the Opal Polaris 7 colour IHC detection kit. DAPI was used as a counter stain. Antibodies can be found in **Supplementary Table ST12**.

### Computational Analysis of Multiplex Immunofluorescence (Leica Cell DIVE) Images

Cell segmentation was performed in QuPath (v0.4.3) according to the QuPath guide to multiplex analysis https://qupath.readthedocs.io/en/stable/docs/tutorials/multiplex_analysis.html). Intensity matrices from each sample were read into RStudio (v4.3.1), processed using sctransform and Seurat using the functions previously described. Proximity analysis was completed using spatula (v0.0.0.9).

### RNAScope of Synovial Tissue

RNAscope was carried out on FFPE RA synovial tissue using the BOND RX Autostainer (Leica). Slides were sectioned at 3 μm and baked at 60 °C for 2 hours. The protocol was developed by combining the Multiplex Fluorescent V2 Assay (ACD) and Opal Multiplex Immunohistochemistry Assay (Akoya Biosciences). BOND Dewax solution (Leica, AR9222) was used for deparaffinisation, and antigen retrieval was carried out using the “ACD HIER 15 min with ER2 (95)” program, followed by the “ACD 15 min Protease” program. *DKK3* was stained using the RNAscope™ LS 2.5 Probe-Hs-DKK3 (415788, ACD) probe and co-stained with VE-Cadherin and CD68, followed by Opal fluorescent dyes (Akoya Biosciences; antibody details found in **Supplementary Table ST12**). RNAscope® LS 2.5 3-plex Positive Control Probe (320868, ACD) and RNAscope® 3-plex LS Multiplex Negative Control Probe (320878, ACD) were used to assess the quality and specificity of staining.

### Bulk RNA sequencing data analysis

Analysis of publicly available bulk RNA sequencing data was performed using previously generated count matrices (HUM0207.v1 - NBDC Human Database(dbcls.jp)) and analysed in RStudio (v4.3.1) using DESeq2 (v1.40.2).

### Peripheral Blood Mononuclear Cells (PBMC) Isolation and Collagen Trans-Endothelial Migration Assay

Adhesion and migration of leukocytes through collagens was assessed using 24 well 3.0 µm pore transwell filters (29442-088, VWR) as previously described^47^. The apical and basal surfaces of transwell filters were coated with 50 µl of Collagen Type I (50 µg/mL; CC050, Sigma-Aldrich), 50 µl of Collagen Type IV (50 µg/mL; C6745-1ml, Sigma-Aldrich) and 50 µl of Collagen Type VI (50 µg/mL; 009-001-108, Rockland) and equilibrated with Endothelial Cell Growth media (C-39210, PromoCell) for 24 hours at 37 °C. Human Umbilical Vein Endothelial Cells (HUVECs; C-12200, PromoCell) were seeded on the apical surface of uncoated or collagen-coated filters for 24 hours at 37 °C before stimulating with TNF-α (100 U/mL; 300-01A-10ug, Sigma-Aldrich) and IFN-γ (10 ng/mL; 285-IF-100/CF, BioTechne) for 24 hours at 37 °C.

PBMCs were isolated from whole blood using SepMate™ PBMC Isolation Tubes (85450, Stem Cell Technologies) according to manufacturer’s protocol. Briefly, whole blood was diluted in an equivalent volume of MACS buffer (PBS + 0.5% BSA + 2mM EDTA) and pipetted gently onto the recommended volume of Lymphoprep™ (85450, Stem Cell Technologies) density gradient medium in SepMate™ tubes and centrifuged at 1200xg for 10 minutes at RT. The enriched PBMC layer was poured into a new tube and washed twice with MACS buffer, centrifuging at 300xg for 8 minutes.

HUVECs were washed to remove residual cytokines, 700 µl fresh Medium 199 (M199; 1544055, Gibco) + 0.15% BSA (A1000801, Life Technologies Ltd) was added into the lower chamber and 200µl PBMCs (2 x 10^6^ cells/mL) in M199 + 0.15% BSA was added to the upper chamber. PBMCs and HUVECs were incubated for 24 hours at 37 °C to allow the cells to adhere and migrate through collagen and HUVECs. Migration was stopped by transferring the filter into a fresh well. Transmigrated leukocytes in the original lower chamber, including those rinsed gently off the basal side of the filter, were collected as the “migrated” fraction; suspended cells in the upper chamber were collected as the “non-adherent” fraction; and adherent cells were detached from the apical surface of the filter using cell dissociation solution (C5914, Sigma-Aldrich) and collected as the “adherent” fraction.

### Flow Cytometric Analysis of Trans-Endothelial Migration Assay

The surface phenotypes of non-adherent (upper chamber), adherent (on filter) and transmigrated (lower chamber) lymphocytes were assessed by flow cytometry. Lymphocytes were labelled with Zombie Aqua viability dye (423102, BioLegend), anti-CD45, anti-CD56, anti-CD3, anti-CD19, anti-CD14, anti-CD4, anti-CD44, anti-CD62L (all from BioLegend) and anti-CD8 (Life Technologies Ltd) for 30 mins on ice. Full antibody details are found in **Supplementary Table ST12**. Sample volume was recorded to calculate absolute cell numbers using the CountBright™ Absolute Counting Beads kit (C36950, ThermoFisher) protocol. Data was acquired on LSRFortessa™ X20 (BD Biosciences) flow cytometer.

Analysis of flow cytometry data was performed in FlowJo (v10.9.0). From the known absolute number of leukocytes added, the percentage of each cell type that adhered and the percentage of each cell types that transmigrated were calculated.

### Sample Preparation, Library Construction, and Sequencing for GeoMx® Digital Spatial Profiler (NanoString)

Synovial tissue FFPE sections from seven RA patient biopsies were selected to assess gene expression in regions of interest (ROIs) using the GeoMx® Digital Spatial Profiler (NanoString). The sections were prepared according to the GeoMX Manual Slide Prep user manual. Briefly, 4 μm sections were cut and baked at 60 °C overnight, deparaffinised, and dehydrated, and antigen retrieval was performed using pH6 citrate buffer. A photocleavable RNA probe-set (Human NGS Whole Transcriptome Atlas RNA:HW0050323, NanoString) was then applied to the sections. The sections were stained for protein detection of CD31, CD90 and CD45, and a nuclear counterstain (Syto Green fluorescent nuclei acid stain, S7575, Thermofisher) applied. A secondary stain to detect the anti-CD31 antibody was used. Full antibody details found in **Supplementary Table ST12.** Slides were loaded in the GeoMx ® Digital Spatial Profiler and ROIs were selected using GeoMx ® Digital Spatial Profiler software, with a CD45 mask applied to exclude capture of the majority of CD45 cells, enriching for capture of fibroblast genes. Endothelial-cell only regions were also selected as a control. RNA probes from each ROI were collected using UV cleaving and stored in separate wells of a 96 well plate. Library preparation and sequencing performed by SourceBioscience (Cambridge, UK), at a sequencing depth of 192 reads/µm^2^ and 5% PHIX.

### Computational Analysis of GeoMx® Digital Spatial Profiler (Nanostring)

Sequencing data was processed using GeoMxNGSPipeline (v2.3.3.10) and read into RStudio (v4.3.1). QC and downstream analysis were performed using NanoStringNCTools (v1.8.0), GeomxTools (3.4.0), and GeoMxWorkflows (1.6.0). ROIs that passed QC were analysed using QuPath (v0.4.3), assessing the number of CD45+ cells in a 30 µm area around the vessel of interest. Regions with an expression of CD31/PECAM below 4.7 were selected, which was below one standard deviation of the mean of CD31/PECAM expression in the endothelial cell positive control regions. Regions were also selected with PTPRC expression below 2.10 to exclude ROIs with significant CD45+ cell inclusion. Endothelium and intra-vessel areas were excluded from the count and values generated of CD45+ cells/area per vessel. Additional vessels present within the broader ROIs were also excluded from the area count. Gene expression from the sequencing data was assessed compared to CD45+ cells/area values taken from the QuPath analysis.

### Animal Models, Preparation of Mouse Synovial Tissue for Single Cell RNA Sequencing (10x Genomics) and Computational Analysis

All animal experiments were approved by the UK Home Office and conducted in accordance with the UK’s Animals (Scientific Procedures) Act 1986 and the UK Home Office Code of Practice. The project and experimental protocols were approved by the University of Birmingham Animal Ethics Review Committee who provided ethical oversight of the study. C57BL/6 mice were purchased from Charles River. All mice used in experimental studies were males aged 8-10 weeks old. Single animals were considered as experimental units.

All mice used for methylated bovine serum albumin (mBSA) antigen-induced arthritis were male, wild type C57BL/6, aged 6-10 weeks. On day −21, mice were inoculated on either lateral side of the lower spine with two 50 µL subcutaneous injections containing emulsified incomplete Freund’s adjuvant (F5506, Sigma Aldrich) supplemented with Mycobacterium tuberculosis H37Ra (2 mg/mL 231141, BD), and mBSA (1 mg/mL, A1009-1G, Sigma Aldrich). For arthritis induction, on day 0, mice received an intra-articular injection directly into a single knee joint containing 10 µL of mBSA (10 mg/mL) in PBS. Arthritis swelling was measured daily between day 0-14 using callipers. Bones with intact joints from hind limbs of day 0 (resting), day 2 (peak), day 7 (resolving) or day 14 (resolved) (*n* = 3 biological replicate samples, each consisted of cells isolated from the joints of three animals) were dissected and transferred into RPMI-1640 (21870084, Gibco™) + 2% FCS containing collagenase D (0.1 g/ml; 11088866001, Roche) and Deoxyribonuclease I (DNase I) (0.01 g/ml; DN25-100mg, Sigma-Aldrich). Samples were incubated at 37 °C for 45 minutes, followed by incubation with medium containing collagenase dispase (0.1 g/ml; 11097113001, Roche) and DNase I (0.01 g/ml) at 37 °C for 30 minutes. Live CD45− and CD45+ (Biolegend, 30-F11) synovial cells were sort purified using the FACSAria (BD BioSciences) and captured with the 10x Genomics Chromium system. Sequencing libraries were generated using the 10x Genomics Single Cell 3′ Solution (version 2) kit and subjected to Illumina sequencing. Alignment to mm10 was completed using CellRanger (10x Genomics, v5). Analysis was completed using RStudio (v4.3.1) and Seurat (v4.3.0). The following Seurat functions were used to process murine data: NormalizeData(), FindVaribleFeatures(), ScaleData(), RunPCA(), RunUMAP(), FindNeighbours(), FindClusters(). Samples were integrated using FindIntegrationAnchors() and IntegrateData(). Data were harmonised (where required) using Harmony (v0.1). Volcano plots were plotted using EnhancedVolcano (v1.13.2) and heatmaps were generated using ComplexHeatmap (v2.10) R packages.

### Organoid Generation, Immunofluorescence Staining and Quantification

HUVECs (C-12200, PromoCell) were cultured in Endothelial Cell Growth Medium (C-39210, PromoCell) according to recommended protocols. Organoids containing synovial fibroblasts only or synovial fibroblasts and HUVECs were generated as previously described^12^. Briefly, 200,000 fibroblasts or 200,000 fibroblasts + 200,000 HUVECs were resuspended in a droplet (35 µL) of Matrigel (356231, Corning) and seeded into polyHEMA (P3932, Sigma-Aldrich)-coated cell culture plates for 21 days.

Following the end of the culture period, organoids were washed in PBS and fixed for 30 minutes at room temperature in ice-cold 4% PFA (43368.9M, ThermoFisher). Following fixation, organoids were washed and then subject to dehydration by sequential immersion in 70% ethanol, 80% ethanol, 100% ethanol (x2) and Histoclear (x3) for 15 minutes each before embedding in paraffin.

FFPE organoid sections were deparaffinised and rehydrated, and antigen retrieval performed at 96°C for one hour using pH9 Dako Target Retrieval Solution (S236784-2, Agilent). Sections were stained for CD31 and either collagen I, collagen VI or POSTN. Secondary antibodies were used to detect the anti-CD31 and anti-POSTN antibodies, whereas anti-collagen I and anti-collagen IV were conjugated to Alexa Fluor 488. Full antibody details can be found in **Supplementary Table ST12**. Following staining, sections were incubated with Vector® TrueVIEW® Autofluorescence Quenching Kit (SP-8400, Vector Laboratories) for 3 minutes. After washing, slides were mounted using VECTASHIELD Vibrance Antifade Mounting Medium with DAPI (H-1800-2, Vector Laboratories).

Whole organoid immunofluorescence images were taken using a Axio Scan Z1 Slide Scanner (Zeiss). For region of interest (ROI) selection, elliptical areas were selected 445 µm^2^ (+/- 5 µm^2^) in size, either 5-10 µm (“close”) or >25 µm (“far”) from endothelial cells, with the marker of interest’s channel turned off. Image analysis was performed using QuPath (v0.4.3).

## Data Availability

Raw counts for the human fibroblast atlas were kindly sent from Prof. Korsunsky and are publicly available at: https://sandbox.zenodo.org/record/772596#.YGsi6BRKg-Q. Processed scRNA-seq data from Alivernini et al., 2020^24^ were kindly provided by Prof. Kurowska-Stolarska and are publicly available at: E-MTAB-8322, E-MTAB-8873 and E-MTAB-8316. Counts from cytokine stimulated human RA fibroblasts (Hum0207)^22^ were obtained from HUM0207.v1 - NBDC Human Database (dbcls.jp). Raw scRNA-seq counts from Wei et al., 2020^12^ were kindly provided by Prof. Korsunsky and is publicly available from Immport (SDY1599) and GSE145286. ScRNA-seq data from Zhang et al., 2019^8^ was obtained from Immport (SDY998). Raw count scRNA-seq data from Zhang et al., 2023^13^ was kindly provided by Prof. Wei and is publicly available from Deconstruction of rheumatoid arthritis synovium defines inflammatory subtypes - syn52297840 - Datasets (synapse.org). Data from Tang et al., 2024^25^ is available at GSE216651. All newly generated sequencing data from this paper will be made publicly available upon publication.

## Code Availability

All code used to process and plot data can be found at chrismahony/Nisa_Mahony_et_al_CroftLab (github.com) and will be made publicly available upon publication.

## Acknowledgements

We thank Genomics Birmingham, Birmingham Tissue Analytics, Biomedical Services Unit and Microscopy Facility at the University of Birmingham and the Digital Pathology Omics Core (DPOC) at the Kennedy Institute for Rheumatology, University of Oxford for technical support.

## Author Contributions

Design of the study: P.N., C.M., A.P.C.; acquisition of data: P.N., A.H., E.C., P.S.C, L-J.M., S.K., C.G.S., and G.N.; analysis and interpretation of data: P.N., C.M., E.C., P.S.C., L-J.M., C.B., S.K., J.D.T., and J.B.R.; provision of human samples, clinical data and histological analysis: R.S., A.F., and K.R.; experimental design advice and access to facilities: K.S.M., H.M.M., M.C., and C.D.B.; provision of scRNA-seq data: K.W.; preparation of the manuscript: P.N., C.M., A.H., and A.P.C. All authors discussed the results and commented on the manuscript.

---

**Figure S1.**
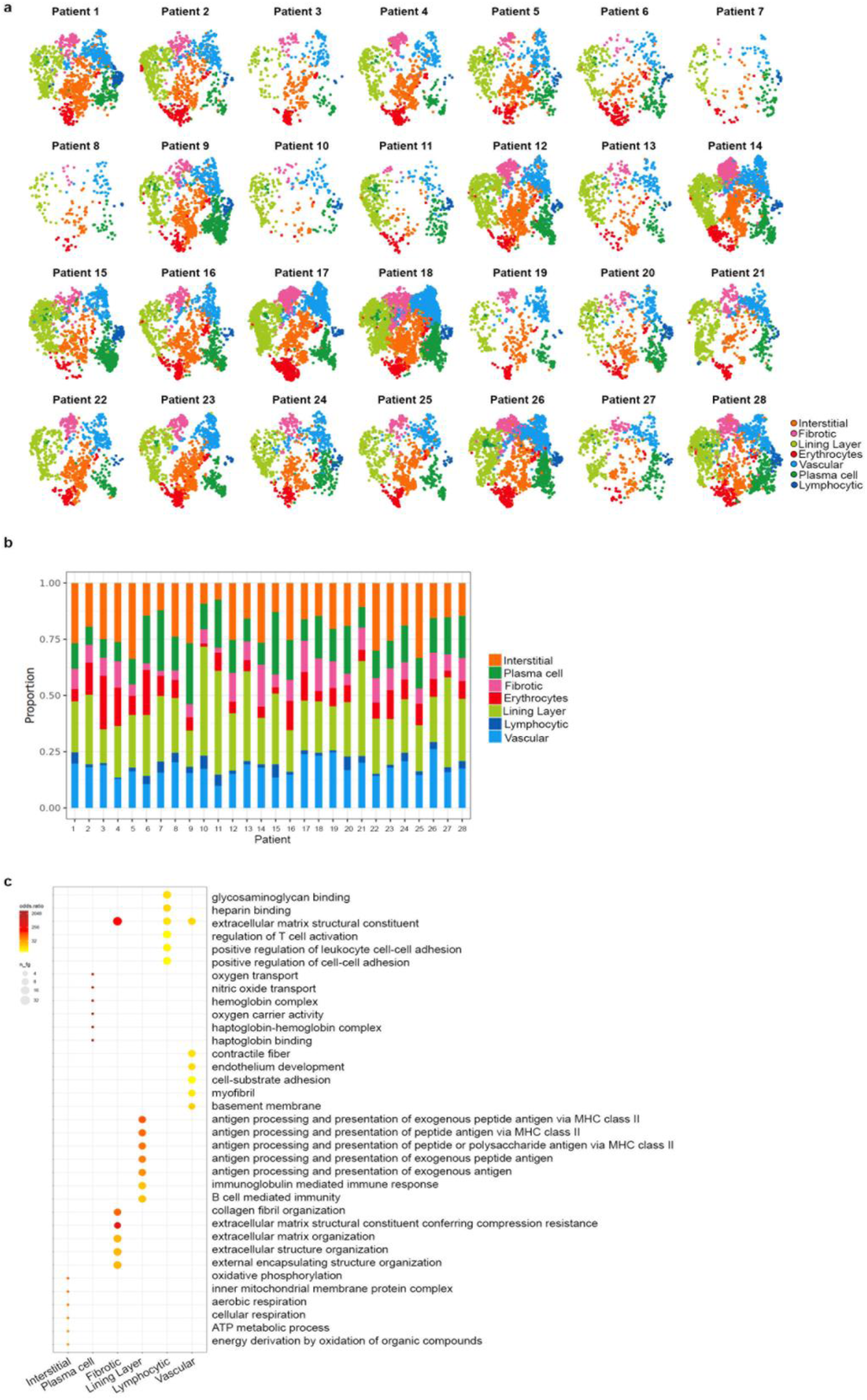
Niche variation per sample. **a**) UMAPs of each ST sample coloured by niche annotation. **b**) Proportion plot of each niche per sample. **c**) GO terms for each niche.

**Figure S2.**
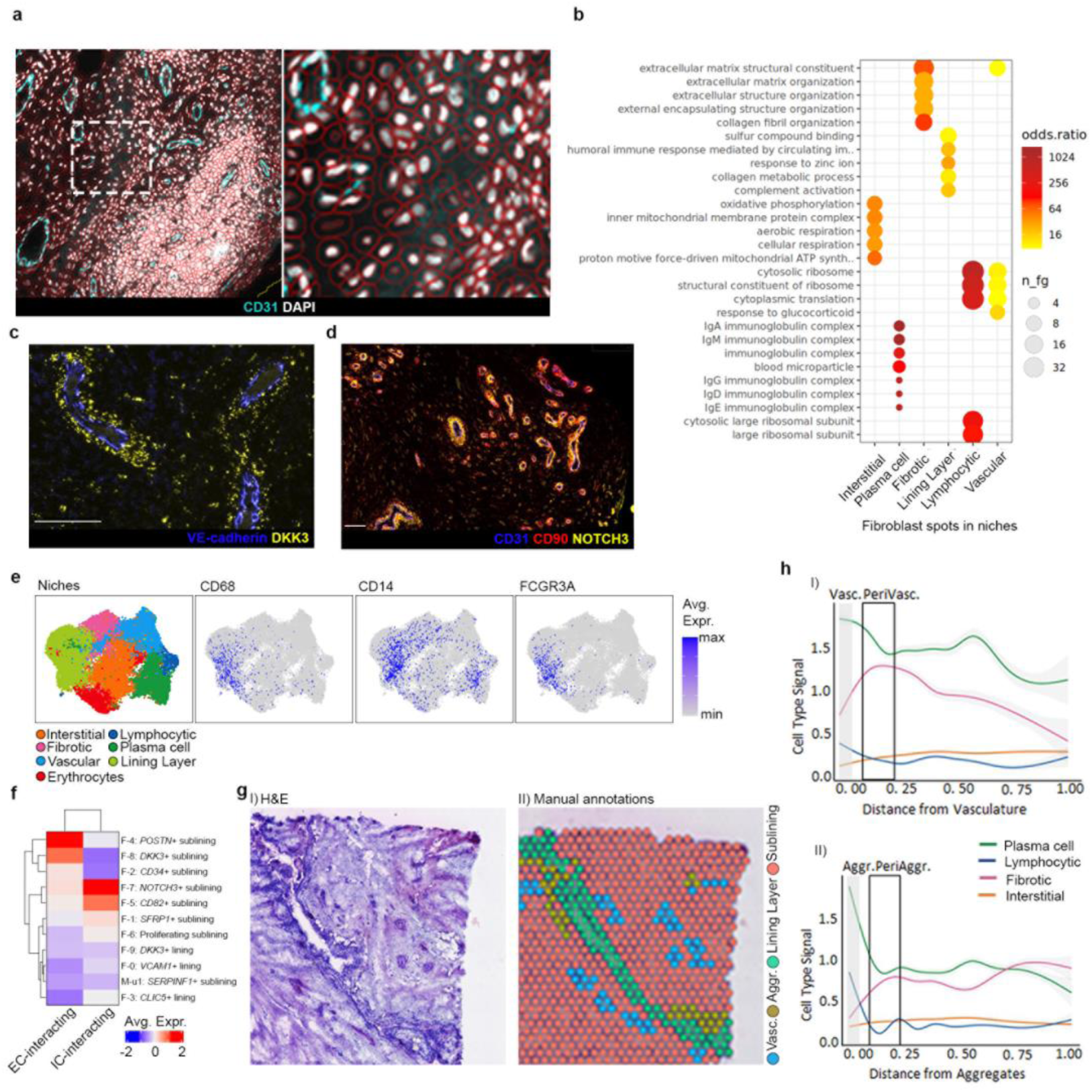
Spatial niche validation. **a**) Representative image showing cell segmentation in QuPath. **b**) GO term analysis of fibroblast-rich spots in each niche. **c**) RNAScope demonstrating perivascular pattern of *DKK3* expression using RNA probe. **d**) Multiplex IF staining showing NOTCH3 perivascular expression. **e**) Feature plots showing expression of macrophage genes in ST. **f**) Heatmap showing expression of EC- and IC-interacting genes in fibroblast clusters from Zhang et al., 2023. **g**) Haematoxylin and eosin staining (I) and ST spatial plot (II) showing manual annotations. **h**) Distance analysis showing changes in niche signal over distance from vasculature (I) and aggregates (II).

**Figure S3.**
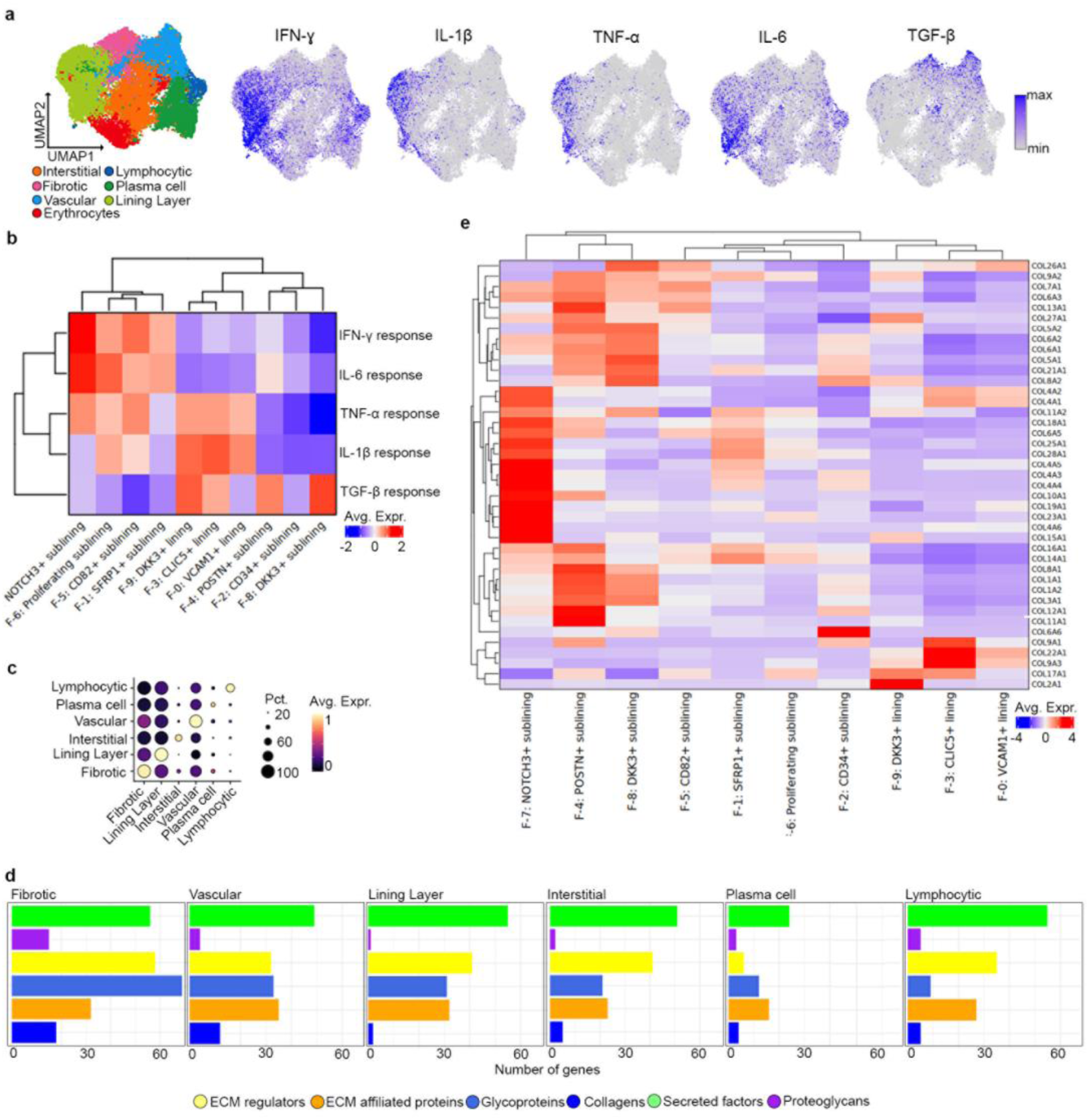
Cytokine and ECM gene program expression in fibroblasts. **a**) Feature plots of expression of fibroblast cytokine response gene modules. Genes were defined as upregulated genes (adjusted p value < 0.05 and FC > 2) with each cytokine stimulation in RA fibroblasts compared to unstimulated control (hum0207). **b**) Expression of cytokine response genes in fibroblast clusters from Zhang et al., 2023. **c)** Dotplot of niche-specific matrisome module in each ST niche. **d**) Number of ECM-related genes categorised by matrisome category in each ST niche. **e**) Expression of collagen genes in fibroblast clusters from Zhang et al., 2023.

**Figure S4.**
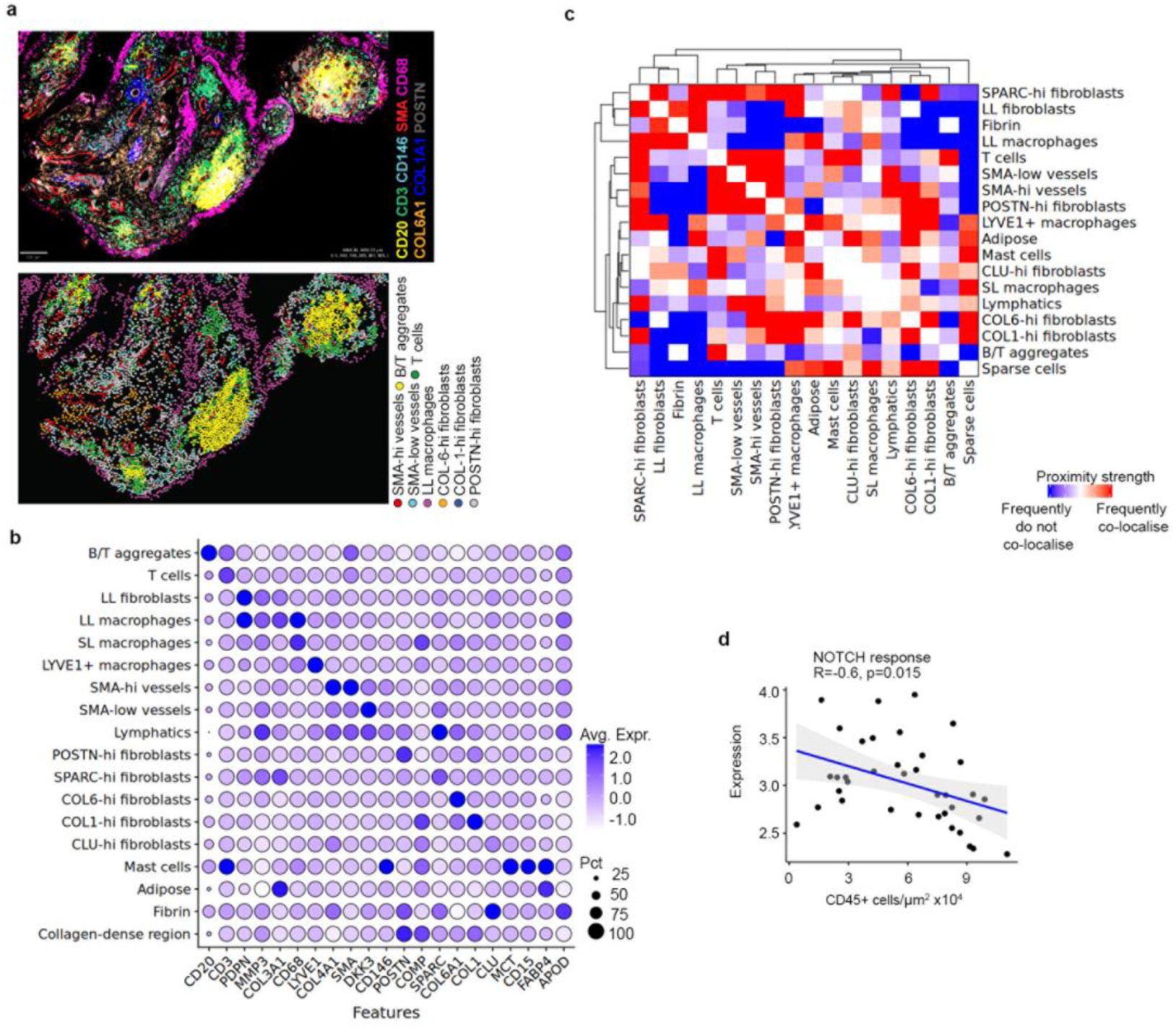
Annotation of cell types in synovial tissue using multiplex IF. **a)** Cell DIVE (Leica) annotation of cell types based on marker genes. **b)** Dotplot showing expression of marker genes across annotated populations. **c)** Proximity analysis of all annotated populations. **d)** Correlation between expression of NOTCH response genes and number of CD45+ cells x10^4^/µm^2^ within each region of interest of synovial tissue samples in GeoMx® Digital Spatial Profiler (NanoString) data. P value calculated using Pearson’s correlation in ggpubr. NOTCH response gene defined by top 40 genes enriched in synovial organoids composed of fibroblasts cocultured with HUVECs compared to fibroblast only-organoids and fibroblasts with HUVECs and DAPT treatment from (Wei et al., 2020).

**Figure S5.**
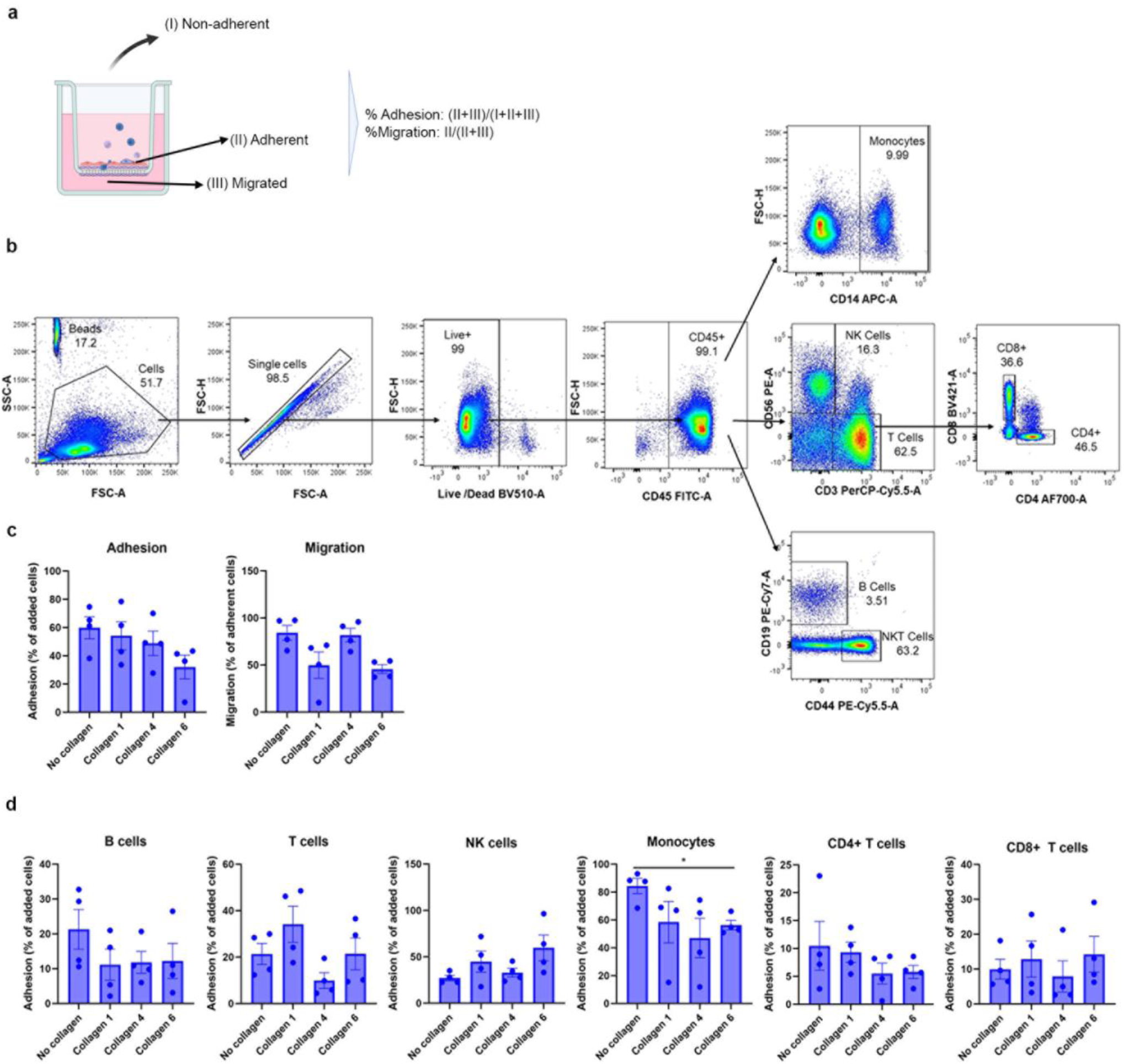
Transwell migration assay. **a**) Schematic overview of leukocyte transwell migration assay. **b**) Flow cytometry gating strategy. **c**) Quantification of adhesion and migration of total cells from flow cytometry analysis. Data is mean ± SEM. **d**) Quantification of adhesion of cells, by cell type, from flow cytometry analysis. Data is mean ± SEM. N=3 for each condition, from induvial donors. **e)** Quantification of migration of cells, split by cell type, from flow cytometry analysis. *, p < 0.05; **, p < 0.01 (determined by Brown-Forsythe ONE-way ANOVA test with Dunnett’s T3 multiple comparisons test).

**Figure S6.**
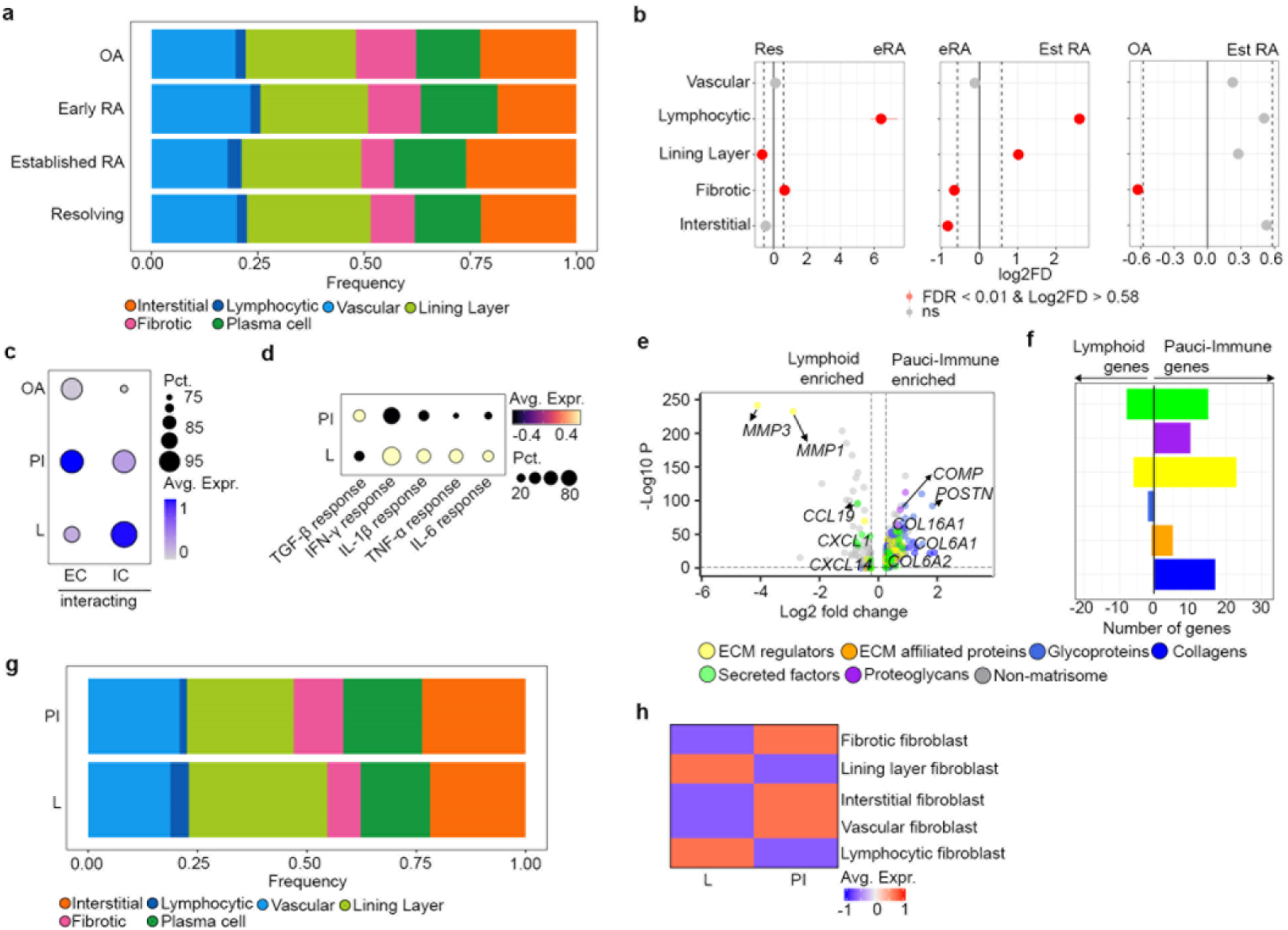
Analysis of fibroblast niches across arthritis disease groups and pathotypes. **a)** Proportion of niches in ST data in early RA, resolving arthritis, established RA and OA. **b)** Relative niche abundance from multiplex IF between resolving arthritis and early RA; established RA and early RA; and established RA and OA samples. **c)** Expression of a module of genes defining IC-interacting fibroblasts and EC-interacting fibroblasts across in fibroblast-rich spots between pauci-immune (PI), lymphoid (L), and osteoarthritis (OA) synovial tissue. **d)** Dotplot of average scaled expression of cytokine response gene modules by fibroblast-rich spots across all cells in lymphoid and PI synovial tissue. **e)** Volcano plot showing differential expression of ECM-related genes in fibroblast-rich spots between lymphoid and PI samples, coloured by Matrisome category. **f)** Number of differentially expressed ECM-related genes in fibroblast-rich spots between lymphoid and PI samples, coloured by Matrisome category. **g)** Proportion of niches between PI and lymphoid tissues in ST data. **h)** Heatmap of expression of genes in fibroblast-rich spots between lymphoid and PI pathotypes of synovial tissue in our ST data.

**Figure S7.**
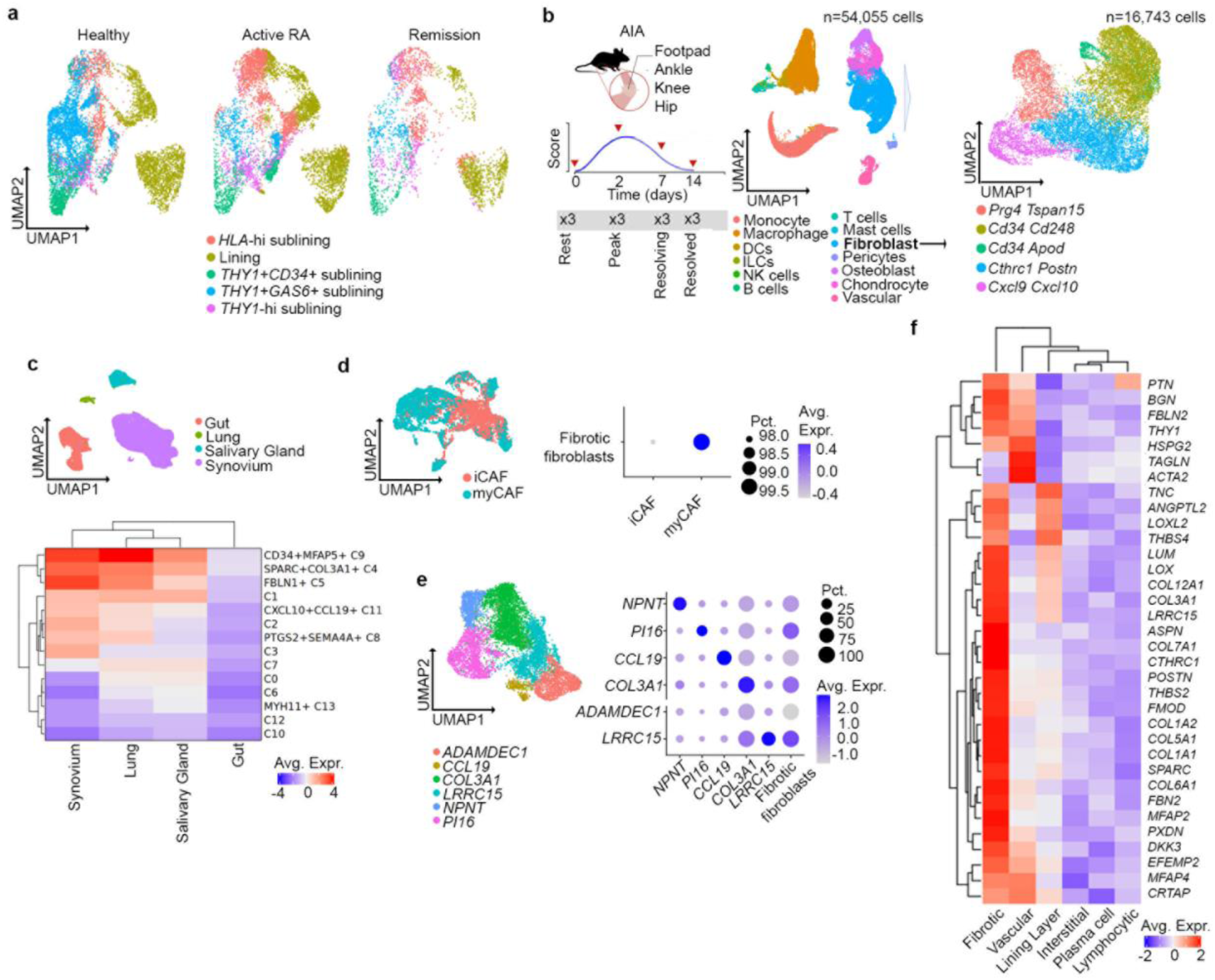
Analysis of fibroblast niches across datasets. **a)** UMAP projection of healthy, active RA and remission synovial tissue samples (Alivernini et al. 2020; Faust et al. 2024). **b)** Overview of antigen-induced arthritis (AIA) timecourse mouse model data, UMAP projection of all clusters and fibroblast sub-clusters. **c)** UMAP of inflammatory diseases from (Korsunsky et al., 2022) and heatmap of the fibrotic niche fibroblast gene markers in 4 tissues analysed from the cross-tissue atlas. **d)** UMAP of cancer-associated fibroblasts signature in breast cancer from GSE176078 (Wu et al., 2021) and dotplot showing expression of fibrotic niche fibroblast gene markers in CAFs. **e)** UMAP of fibroblast subsets of Buechler et al., 2021 and dotplot of expression of identified CAF cluster alongside fibrotic niche fibroblast gene program expression. **f)** Heatmap of gene expression of myofibroblast gene expression profiles in fibroblast niche from our ST data.

